# Essential role of Hepcidin in host resistance to disseminated candidiasis

**DOI:** 10.1101/2024.10.29.620511

**Authors:** Tanmay Arekar, Divya Katikaneni, Sadat Kasem, Dhruv Desai, Thrisha Acharya, Augustina Cole, Nazli Khodayari, Sophie Vaulont, Bernhard Hube, Elizabeta Nemeth, Alexander Drakesmith, Michail S Lionakis, Borna Mehrad, Yogesh Scindia

## Abstract

*Candida albicans* is a leading cause of life-threatening invasive infections with up to 40% mortality rates in hospitalized individuals despite antifungal therapy. Patients with chronic liver disease are at an increased risk of candidemia, but the mechanisms underlying this susceptibility are incompletely defined. One consequence of chronic liver disease is attenuated ability to produce hepcidin and maintain organismal control of iron homeostasis. To address the biology underlying this critical clinical problem, we demonstrate the mechanistic link between hepcidin insufficiency and candida infection using genetic and inducible hepcidin knockout mice. Hepcidin deficiency led to unrestrained fungal growth and increased transition to the invasive hypha morphology with exposed 1,3, β-glucan that exacerbated kidney injury, independent of the fungal pore-forming toxin candidalysin in immunocompetent mice. Of translational relevance, the therapeutic administration of PR-73, a hepcidin mimetic, improved the outcomes of infection. Thus, we identify hepcidin deficiency as a novel host susceptibility factor against *C. albicans* and hepcidin mimetics as a potential intervention.

## INTRODUCTION

*Candida albicans* is a commensal fungus and, in >80% of the population, is a part of the normal flora of the gut microbiota^1,2^. Its transition from commensalism to pathogenicity remains the predominant cause of invasive candidiasis ^3–7^, with up to 40% mortality despite antifungal therapy ^8^. For incompletely understood reasons, the kidneys are a major target of infection, as evidenced by autopsies of patients with disseminated candidiasis^9–11^ and in animal models^12–17^. For patients with a high risk of dissemination, The Infectious Diseases Society of America guidelines recommend imaging the kidneys to exclude abscesses, fungal balls, or urologic abnormality^18^. However, while the risk of systemic candidiasis and clinical outcomes vary significantly^19,20^, host-related risk factors and associated mechanisms are incompletely defined. Thus, host susceptibility factors that promote or limit fungal growth must be identified to complement the current complement of antifungal drugs.

Iron is an essential micronutrient for host physiology^21,22^ as well as a nutrient and virulence factor for *C. albicans*^23–31^. Iron supplementation increases the resistance of *C. albicans* to antifungal agents^32^. Injecting mice with iron preparations worsens outcomes^33,34^ and impaired iron acquisition of *C. albicans* reduces the capacity to damage epithelial cells^35^. Collectively, these studies identify iron as a crucial component for *C. albicans*^28,36^ and support the notion that a dysregulated host iron metabolism may increase susceptibility to disseminated candidiasis. However, the current understanding of host iron content as a susceptibility factor to candida infections is based on supraphysiological injections of iron preparations in animals, thus lacking clinical relevance.

Systemic iron metabolism is regulated primarily by the hepcidin-ferroportin axis. Hepcidin binds to and triggers the internalization of ferroportin, a transmembrane protein that transfers intracellular iron to the plasma^37,38^. As a part of the evolutionarily conserved nutritional immunity^39^, hepcidin-mediated ferroportin degradation attenuates the rate of entry of iron into plasma and extracellular fluid^40,41^, allowing transferrin to remove non-transferrin-bound iron (NTBI), an iron species that is highly bioavailable to microbes and stimulates their rapid growth^42–44^. The *HAMP* gene encodes hepcidin and is mainly produced by hepatocytes^40,45,46^. While hepcidin production is exquisitely sensitive to iron levels, *HAMP* transcription is also induced by microbial sensing by pattern recognition receptors (PRR) or cytokines like IL-1β, IL- 6, or IL-22^45,47,48^ in a STAT3-dependent manner^47,49,50^.

Hepatic hepcidin levels increase after intravenous infection with *C. albicans*^47^, suggesting that the host activates an iron-withholding program during systemic candidiasis. Candidiasis is the most common fungal infection in chronic liver disease patients and those undergoing liver transplants ^51–54^. However, the mechanisms underlying this increased susceptibility are incompletely defined.

In this study, we identify the essential role of hepcidin in host defense against *C. albicans* in immunocompetent mice. We demonstrate the ability of iron to change fungal wall composition and exacerbate inflammation in hepcidin-deficient mice infected with a candidalysin-deficient *C. albicans* strain that cannot elicit a response in wild-type littermates. We uncover hepcidin-induced iron sequestration as an essential host defense mechanism against the fungal pathogen *C. albicans and* show that treatment of susceptible mice with a hepcidin agonist attenuates fungus-induced kidney pathology.

## RESULTS

### Clinical Implications: Liver fibrosis attenuates hepcidin production and is associated with renal iron overload

To access the translational relevance of our study, we measured the hepatic hepcidin transcripts and serum hepcidin levels in patients with liver diseases (including alpha-1 antitrypsin disease, alcohol-induced cirrhosis, and cirrhosis due to hepatitis B and C virus infection). Compared to control individuals, liver hepcidin transcripts and serum hepcidin levels were significantly lower in patients with chronic liver disease (**Fig. 1A-B**). We then asked whether liver disease-induced loss of hepcidin changes tissue iron distribution. C57BL/6 mice injected with carbon tetrachloride (CCl_4_) developed severe liver fibrosis (**Fig. 1C-D**), and this was associated with attenuated liver *Hamp* gene expression (**Fig. 1E)** and intra-cellular iron deposition in the kidneys (**Fig. 1F-G)**. Given the conserved role of hepcidin in humans and mice, we anticipate that patients with chronic liver disease may accumulate iron in their kidneys, rendering them susceptible to bloodstream candida infections.

**Figure. 1:**
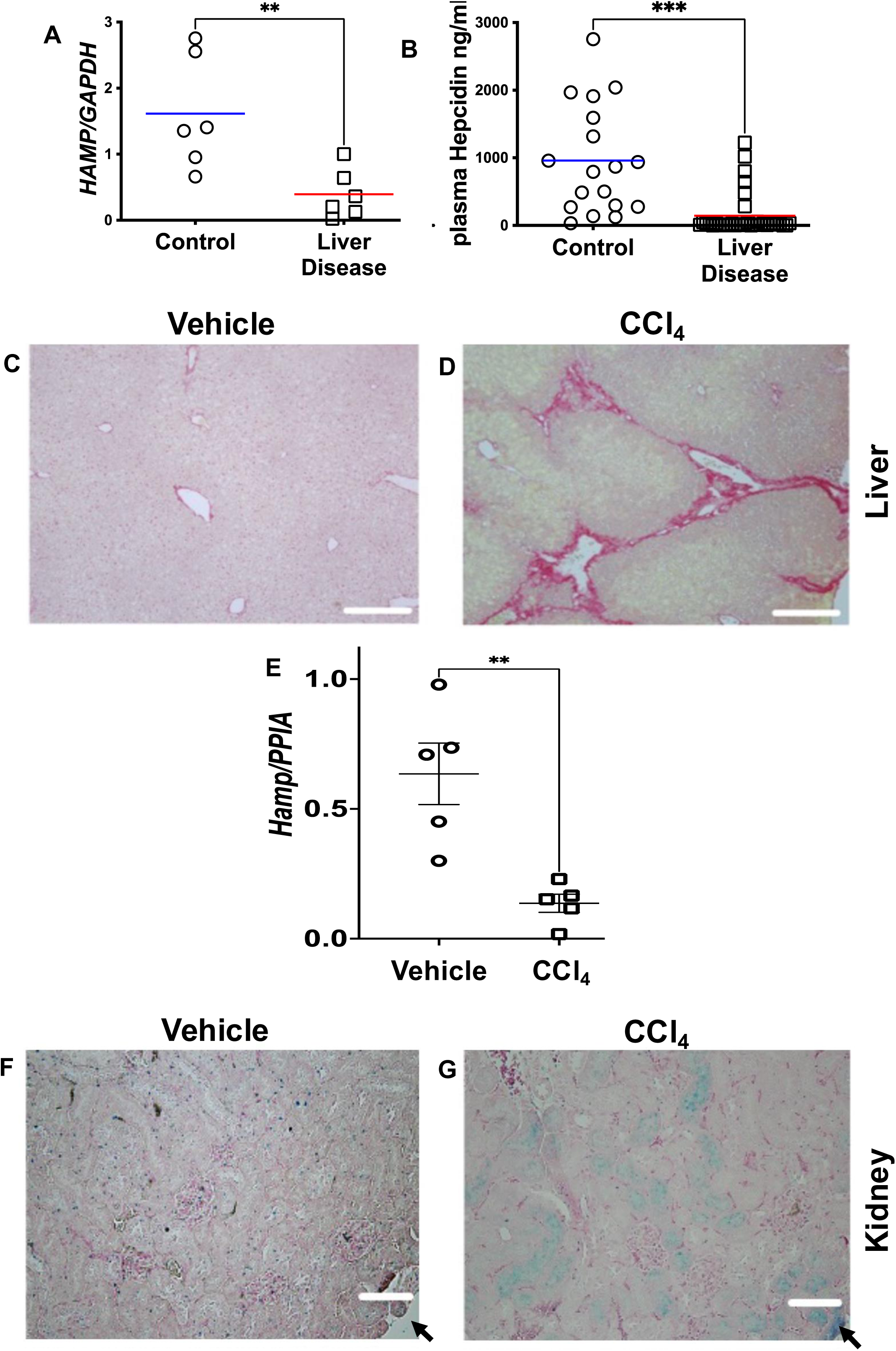
Chronic liver disease attenuates hepcidin production and is associated with renal iron accumulation: Implications for human candidiasis and disease. Compared to patients without liver disease, hepatic hepcidin gene expression and serum hepcidin levels are attenuated in patients with chronic liver disease (**A-B**). 8-10 weeks-old C57BLK/6 mice were injected intraperitoneally with 0.3% CCl_4_ or vehicle (corn oil) at a dose of 10 uL/g of body weight twice weekly for five weeks and euthanized. Compared to the vehicle, Picro Sirius red staining of the CCl_4_ injected livers revealed extensive fibrosis with multiple fibrotic septa and divided hepatic lobules (**C-D**). Scale bar: 50 μM. The intra-hepatic hepcidin gene expression was significantly attenuated in fibrotic livers (**E**). The kidneys of the vehicle and CCl_4_ were stained for Perl’s detectable iron deposits. Perl’s detectable iron deposits were not observed in vehicle-treated kidneys (**F**). However, following CCl_4_ treatment, blue Perl’s detectable iron deposits were observed in the renal tubules (**G)** scale bar: 30 μM, arrow denotes the edge of the section. Each point represents a patient. A 2-tailed Mann-Whitney test determined statistical significance and plotted as mean ± SEM. **P < 0.005, ***P < 0.001.

### Hepcidin is required for resistance to *Candida albicans* infection

To determine whether hepcidin deficiency and associated tissue iron overload are a risk factor in disseminated candidiasis, we compared outcomes of infection between *Hamp^-/-^*mice and their littermates (W.T.) controls. We first confirmed the baseline differences in iron homeostasis between W.T. and *Hamp^-/-^* mice. The spleens of W.T. mice accumulated iron in the red pulp regions, which was not observed in *Hamp^-/-^* mice (**Supplementary** Figure 1A-B). In contrast, unlike W.T. kidneys, the kidneys of *Hamp^-/-^* mice were iron-loaded (**Supplementary** Figure 1C-D). Most of the iron was observed in the cortico-medullary junction and extended to the deep cortex. Most of the iron accumulates in the tubular compartment, with little to none in the glomerular or interstitial regions (**Supplementary** Figure 1E). To test the hypothesis that hepcidin deficiency-associated renal iron accumulation worsens the outcome of disseminated candidiasis, W.T. or *Hamp^-/-^*mice were intravenously injected with SC5314 yeast cells. All *Hamp^-/-^*mice were moribund by day 2-3 post-infection, whereas their W.T. littermates reached this endpoint only at day 6 (**Fig. 2A)**. To avoid mortality in *Hamp^-/-^* mice, we reduced the infecting dose by half and followed mouse outcomes (**Fig. 2B)**. Twenty hours post-infection, there was no candidemia in W.T. and *Hamp^-/-^*mice (**Supplementary** Figure 2A-B), which agrees with previous reports^17^. However, 72 hours post-infection, unlike W.T. littermates, *Hamp^-/-^* mice were lethargic and moribund. Compared to W.T. mice, the renal fungal burden was significantly higher in *Hamp^-/-^* mice (**Fig. 2C-E**). While the liver fungal burden was also significantly higher in *Hamp^-/-^* mice, it was 2 log fold lower than in the kidneys (**Fig. 2F, Supplementary** Figure 2D). The fungal burden was significantly higher in the iron-sufficient W.T. spleens, whereas iron-deficient spleens of *Hamp^-/-^* mice were devoid of fungal colonies (**Fig. 2G, Supplementary** Figure 2C). *C. albicans* in the W.T. kidney were predominantly in yeast-pseudohyphae forms (**Fig. 2H**) but had undergone pronounced hyphal transformation in *Hamp^-/-^* mice (**Fig. 2I**), indicating increased virulence^55^. PAS staining did not reveal obvious pathology in the spleen and livers of W.T. and *Hamp^-/-^* mice (**Supplementary** Figure 2E). Renal histopathology showed occasional small foci of inflammation with fungal abscesses in W.T. kidneys (**Supplementary** Figure 2F**, white dotted circle**) and numerous fungal abscesses in *Hamp^-/-^* mice (**Supplementary** Figure 2G**, white dotted circle**), which correlated with the observed renal fungal burden. Together, this demonstrates that hepcidin deficiency increases kidney susceptibility to *C. albicans*.

**Figure. 2:**
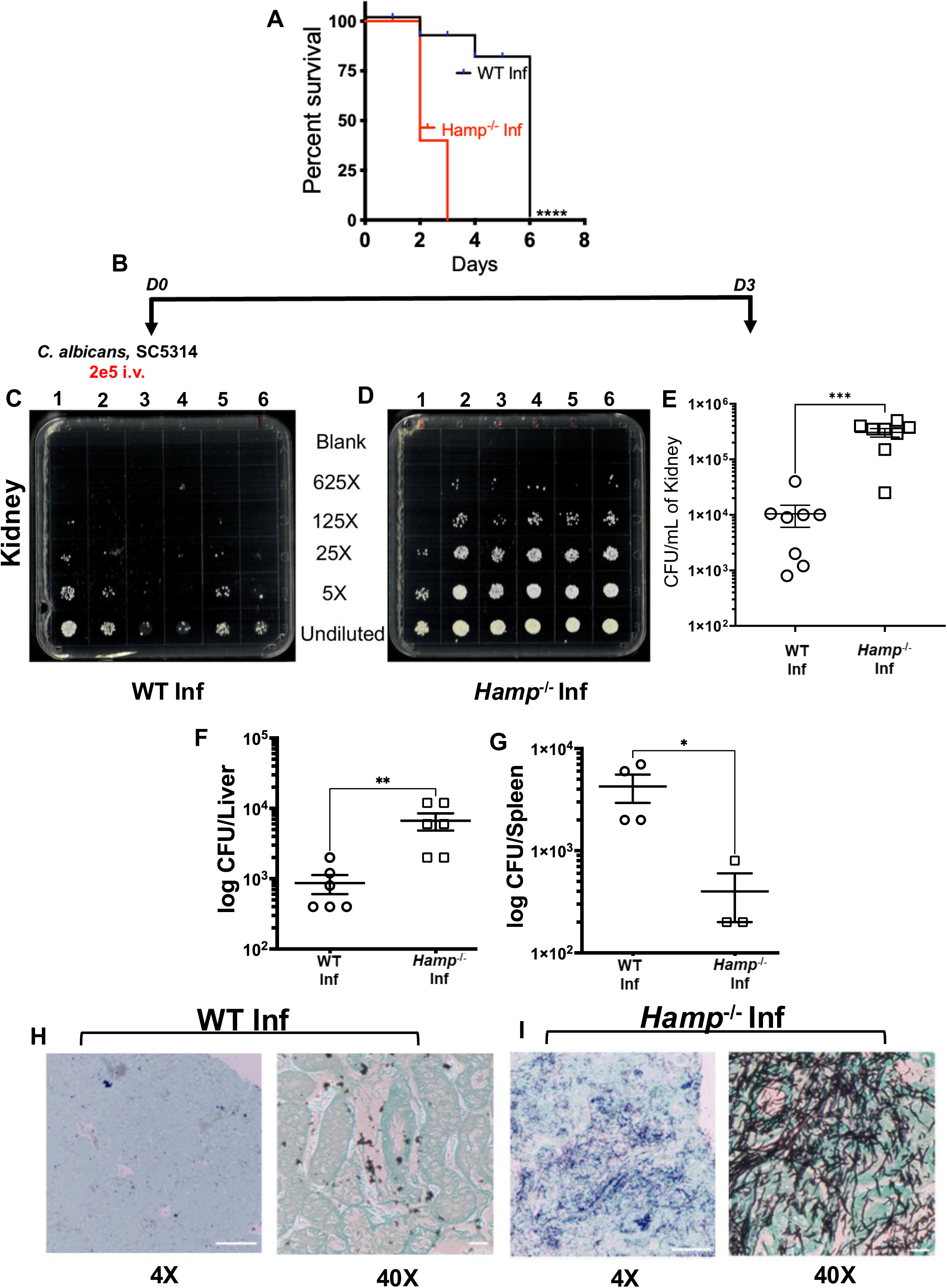
Hepcidin deficiency worsens outcomes of disseminated candidiasis. 12-week-old littermates (W.T.) and *Hamp^-/-^* mice (n= 8 each) were intravenously infected with 4 X 10^5^ *C. albicans* (SC5314). Day 3 post-infection, 10% mortality was observed in W.T. mice, whereas all *Hamp^-/-^*mice died. The W.T. mice reached a similar endpoint by day 6 (**A**). To reduce mortality, mice were intravenously injected with 2 X 10^5^ *C. albicans,* and tissues were harvested three days later (**B**). Compared to W.T. mice, iron-overloaded *Hamp^-/-^* mice have a significantly higher fungal burden in the kidney (**C-E**) and liver (**F**). In contrast, the iron-sufficient W.T. spleens harbored significantly higher fungal colonies than iron-deficient *Hamp^-/-^* spleens (**G**). A 2-tailed Mann-Whitney test determined statistical significance. Data is presented as mean± SEM. *P < 0.05, **P < 0.001, ***P < 0.0001. Grocott methenamine silver staining revealed that the fungus in the W.T. kidney was predominantly in yeast-pseudohyphae form (**H**) but had undergone a hyphal transformation in the *Hamp^-/-^* kidneys (**I**). Scale bar:100 μM and 30 μM.

### Kidney iron provides a local microenvironment for the accelerated growth and virulence of *C. albicans*

Based on the above observations, we evaluated whether the kidney iron content could accelerate the growth of *C. albicans*. Naïve C57BL/6 kidneys were cultured 48 hours with or without ferric ammonium citrate as an iron source, lysed, and inoculated with *C. albicans* yeasts to assess fungal growth. *C. albicans* growth increased linearly with increased iron content of the kidney lysates (**Supplementary** Figure 3A). Since the renal proximal tubular epithelial cells (PTEC) participate in iron recycling^56^ and most of the iron in *Hamp^-/-^* mice accumulates in the renal tubules, we loaded HK-2 cells, a human PTEC cell line, with or without ferric chloride as described previously^57^ (**Supplementary** Figure 3B-C) and exposed them to *C. albicans* for 24 hours. When the culture supernatants from both conditions were spiked with an equal number of *C. albicans* yeast cells, there was significantly higher fungal growth in the supernatant of iron-loaded HK-2 cells (**Supplementary** Figure 3D). To determine whether iron played a role in promoting the virulence of *C. albicans,* the fungus was grown in enriched yeast nitrogen broth [YNB] and supplemented with or without iron as described by Tripathi et al. ^32^ (**Supplementary** Figure 3E and H). Independent of the media’s iron content, under hyphal-inducing temperature (37^0^C), *C. albicans* underwent a yeast-to-hyphae transition in three hours (**Supplementary** Figure 3F and I). However, 24 hours later, only the high iron medium sustained the hyphal morphotype (**Supplementary** Figure 3G and J). Collectively, these data show that the kidney iron content promotes the growth of *C. albicans* and sustains it in its invasive hyphal form.

### Increased fungal burden and hyphal formation in *Hamp^-/-^ mice* worsen acute kidney injury (AKI)

The increased fungal burden and prominent hyphal formation in *Hamp^-/-^*mice were associated with tubular necrosis and increased immune cell infiltration (**Fig. 3A-D).** *C. albicans* infection also significantly compromised renal function, as illustrated by increased plasma creatinine levels, and injured the proximal renal tubules as indicated by increased *Ngal* and *Kim1* expression in the kidneys of *Hamp^-/-^* mice (**Fig. 3E**). Thus, *Hamp^-/-^*mice develop more severe acute kidney injury (AKI) following disseminated candidiasis. The gene expression of intra-renal inflammatory cytokines such as TNFα, IL-1β, and IL-6 positively correlated with the extent of AKI (**Supplementary** Figure 4A-C). To rule out that increased fungal burden and hyphal formation observed in *Hamp^-/-^* mice was due to impaired production of chemoattractants and leukocyte infiltration, we measured the intra-renal gene expression of chemoattractants like *Csf3* (neutrophils), *Ccl2, Cxcl11* (monocytes) and analyzed kidney infiltrating leukocytes by flow cytometry. Uninfected W.T. and *Hamp^-/-^* mice had comparable gene expression of *Csf3, Ccl2,* and *Cxcl11* (**Supplementary** Figure 4D-F). *C. albicans* infection increased the gene expression of all the above chemoattractants in W.T. mice, which were further significantly amplified in *Hamp^-/-^ mice* (**Supplementary** Figure 4D-F). Neutrophil and monocyte numbers in uninfected W.T. and *Hamp^-/-^* kidneys were comparable (**Fig.3H-I, Gating strategy in Supplementary** Figure 5). After infection, neutrophil and monocyte numbers significantly increased in W.T. mice (**Fig. 3H-I**). However, compared to infected W.T. mice, both cell types were significantly increased in infected *Hamp^-/-^*mice (**Fig. 3H-I**).

**Figure. 3:**
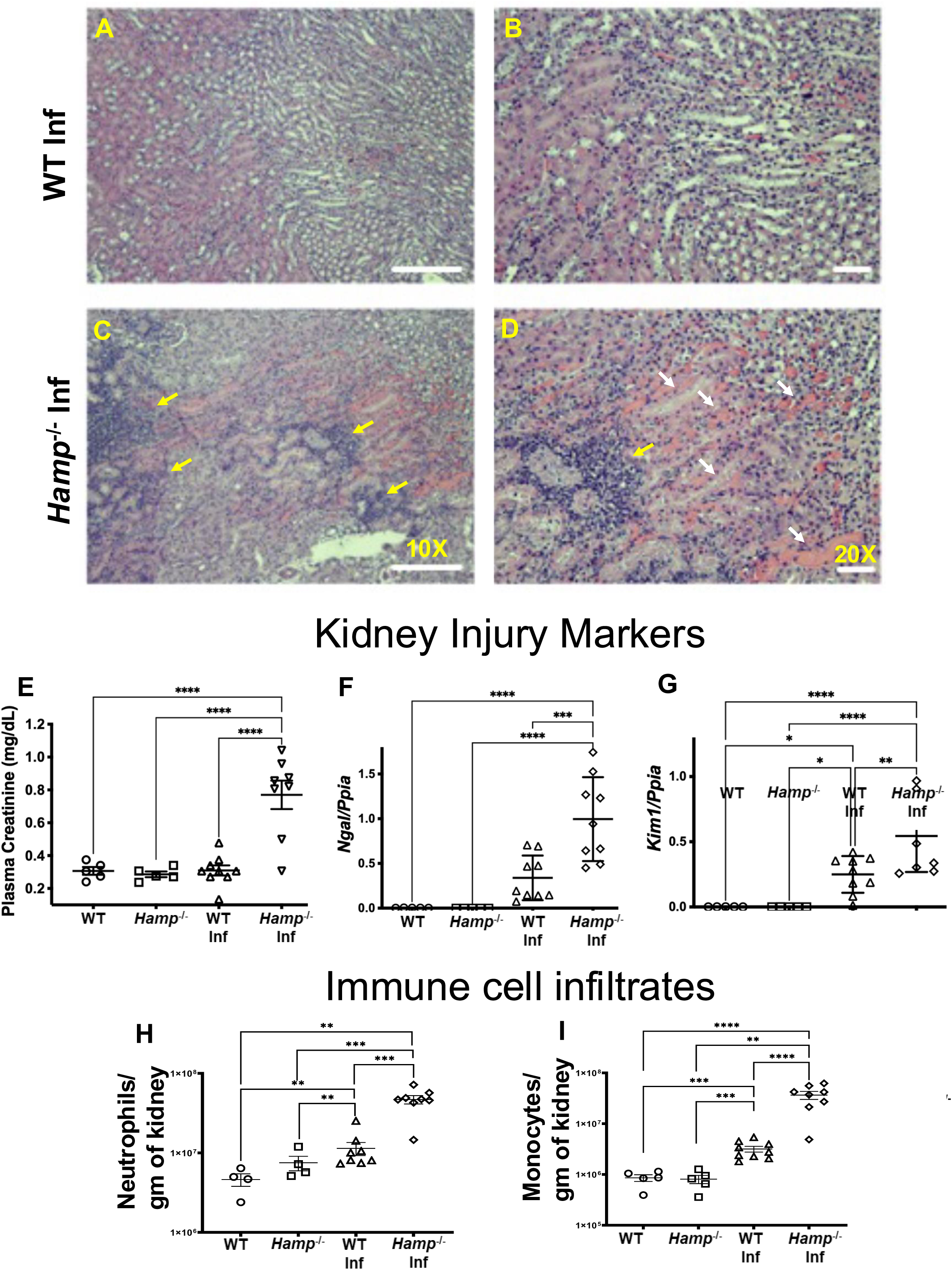
C. albicans exacerbates acute kidney injury (AKI) in hepcidin knockout mice. W.T. and *Hamp^-/-^*mice were infected with 2e^5^ *C. albicans* yeast, and tissues were harvested three days later. Hematoxylin and Eosin (H&E) staining revealed that compared to W.T. kidneys (**A-B**), the *Hamp^-/-^* kidneys had more tubular epithelial cell necrosis (dark pink fragmented cytoplasm with no nuclei), with casts, luminal debris, and multiple foci of inflammation (**C-D**) Scale bar A & C:100 μM, B & D: 50 μM. Markers of kidney injury and inflammation were comparable in naïve W.T. and *Hamp^-/-^*mice. Post infections, AKI, measured by plasma creatinine and proximal tubular injury markers *Ngal* and *Kim1,* was exacerbated in *Hamp^-/-^* mice (**E-G**). The infected *Hamp^-/-^* kidneys had significantly higher numbers of neutrophils and monocytes than W.T. kidneys (**H-I**). Data was analyzed using 2-way ANOVA with Holm-Šídák’s multiple comparisons test and represented as mean ± SEM. *P < 0.05, **P < 0.005, ***P < 0.001. ****P < 0.0001.

### Hyphal transformation in the hepcidin knockout mice is associated with cell death, immune cell infiltration, and loss of kidney parenchyma

Hyphal transformation of *C. albicans* is associated with the expression of *ECE1* (extent of cell elongation 1)^58^, a core filamentation gene expressed under the most hyphal-inducing conditions^59^. *ECE1* codes for the Ece1 polyprotein, the precursor to candidalysin, a pore- forming cytolytic toxin^60,61^. Unlike during W.T. kidney infection, high levels of *ECE1* gene expression was observed in the infected *Hamp^-/-^* kidneys (**Fig. 4A**). Active penetration or induced endocytosis by viable wild-type *C. albicans* hyphae damages epithelial cells via necrosis^62,63^, and interactions with macrophages induce candidalysin-mediated pyroptosis^64,65^. Indeed, in infected *Hamp^-/-^*kidneys, the expression of *ECE1* was associated with increased gene expression of *GsdmD* encoding Gasdermin D, a key factor of pyroptosis, and Mixed lineage kinase domain-like protein (*Mlkl:* Necroptosis), both indicative of inflammatory forms of cell death (**Fig. 4B-C**).

**Figure. 4:**
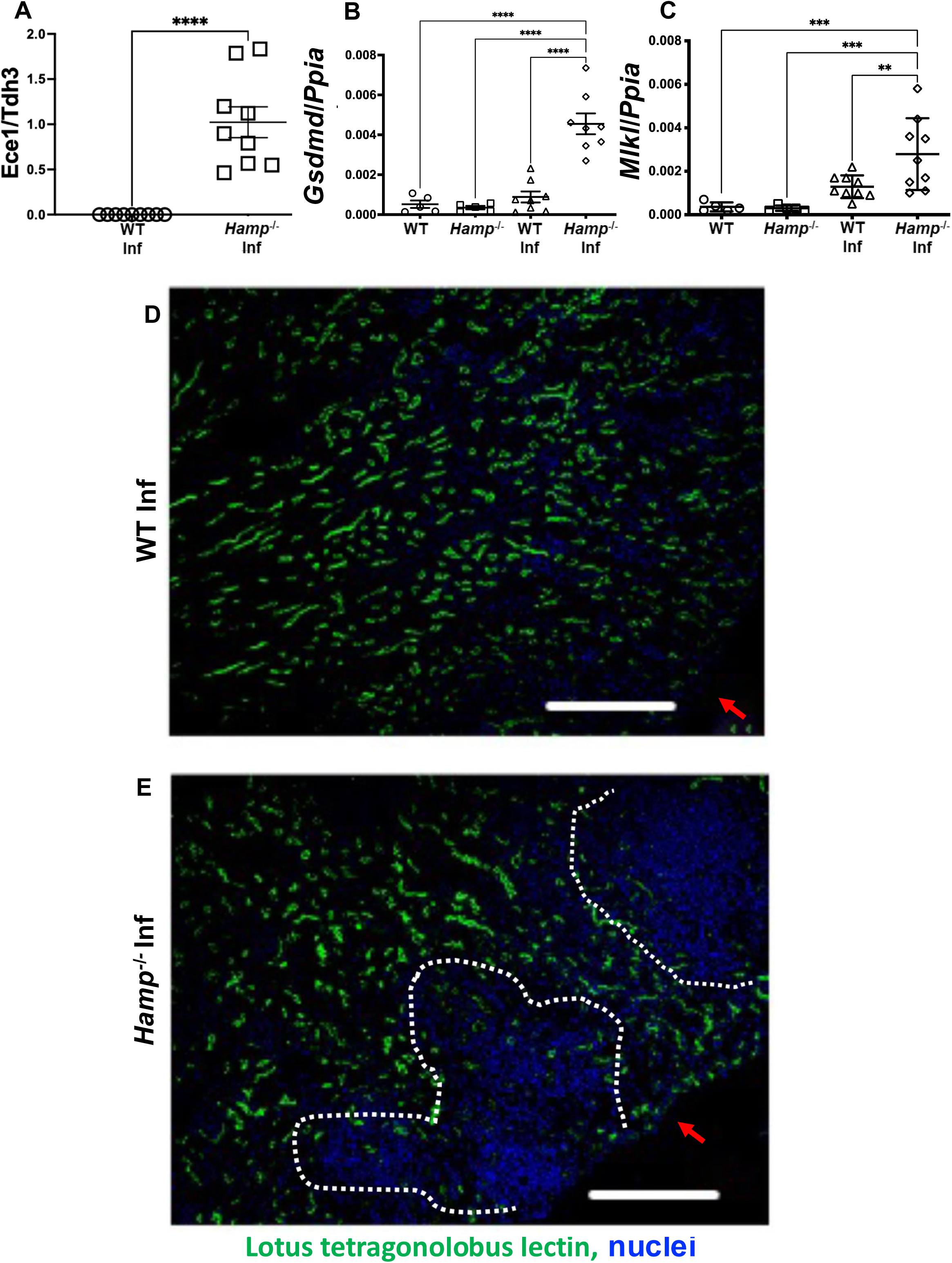
Expression of *Ece1* correlates with cell death, inflammation, and loss of renal parenchyma. 3-days post *C. albicans* infection, the kidneys of W.T. and *Hamp^-/-^* mice were analyzed for expression of the fungal gene *Ece1*. *Ece1* gene expression was observed only in the *Hamp^-/-^* kidneys (**A**). Cell death markers were comparable in naïve W.T. and *Hamp^-/-^* kidneys (**B-C**). *GsdmD* (pyroptosis) and *Mlkl* (necrosis) gene expression were not significantly induced in the infected W.T. kidneys (**B-C**). However, the infected *Hamp^-/-^* kidneys significantly upregulated the expression of *GsdmD* and *Mlkl* (**B-C**). Data was analyzed using 2-way ANOVA with Holm- Šídák’s multiple comparisons test and represented as mean ± SEM. *P < 0.05, **P < 0.005, ***P < 0.001. ****P < 0.0001. The proximal tubular epithelial cells (PTEC) are the most abundant epithelial cells in the kidney. Lotus tetragonolobus lectin (LTL) staining, a marker for PTEC, revealed a uniform distribution throughout the infected W.T. kidneys (**D**) (the red arrow indicates the edge of the section). However, infected *Hamp^-/-^* kidneys presented with multiple sections devoid of LTL-positive PTECs, indicating loss of kidney parenchyma (**E**). Scale bar D & E:100 μM.

Immunofluorescence staining of proximal tubular epithelial cells (PTEC) revealed an even distribution of PTEC in the infected W.T. kidneys (**Fig. 4D**). However, large areas of infected *Hamp^-/-^* kidneys were devoid of PTEC (**Fig. 4E, White dotted circle**), indicating loss of renal parenchyma.

### Hepcidin deficiency in immunocompetent mice induces AKI independent of candidalysin

To delineate the contribution of excessive fungal burden and examine a mechanism for pathology, we first examined the impact of candidalysin. We infected W.T. and *Hamp^-/-^* mice with *C. albicans* lacking the ability to produce candidalysin (*ECE1* null mutants: *ece1*Δ/Δ)^61,66^. Compared to W.T. mice, the *ece1*Δ/Δ strain proliferated significantly more and underwent hyphal transformation in *Hamp^-/-^*mice. This was associated with significantly greater renal injury and immune cell infiltration (**Fig. 5A-H**). While the fungal burden and hyphae formation in *Hamp^-/-^* mice infected with either *ece1*Δ/Δ or an *ece1*Δ revertant strain (*<u>ece1</u>*<u>ΔΔ*+ECE1*</u>, with similar virulence to SC5314)^61,67^ was comparable (**Fig. 5B**), the *ece1*Δ/Δ*strain-*induced significantly less renal injury and inflammation (Fig. **5E-H**), indicating that candidalysin dictates the extent of kidney pathology. However, unlike published literature on leukopenic mice infected with *ece1*Δ/Δ strain^68^, our data identify that hepcidin deficiency in an immunocompetent animal still increases fungal burden, hyphal formation, and kidney pathology, independent of candidalysin.

**Figure. 5:**
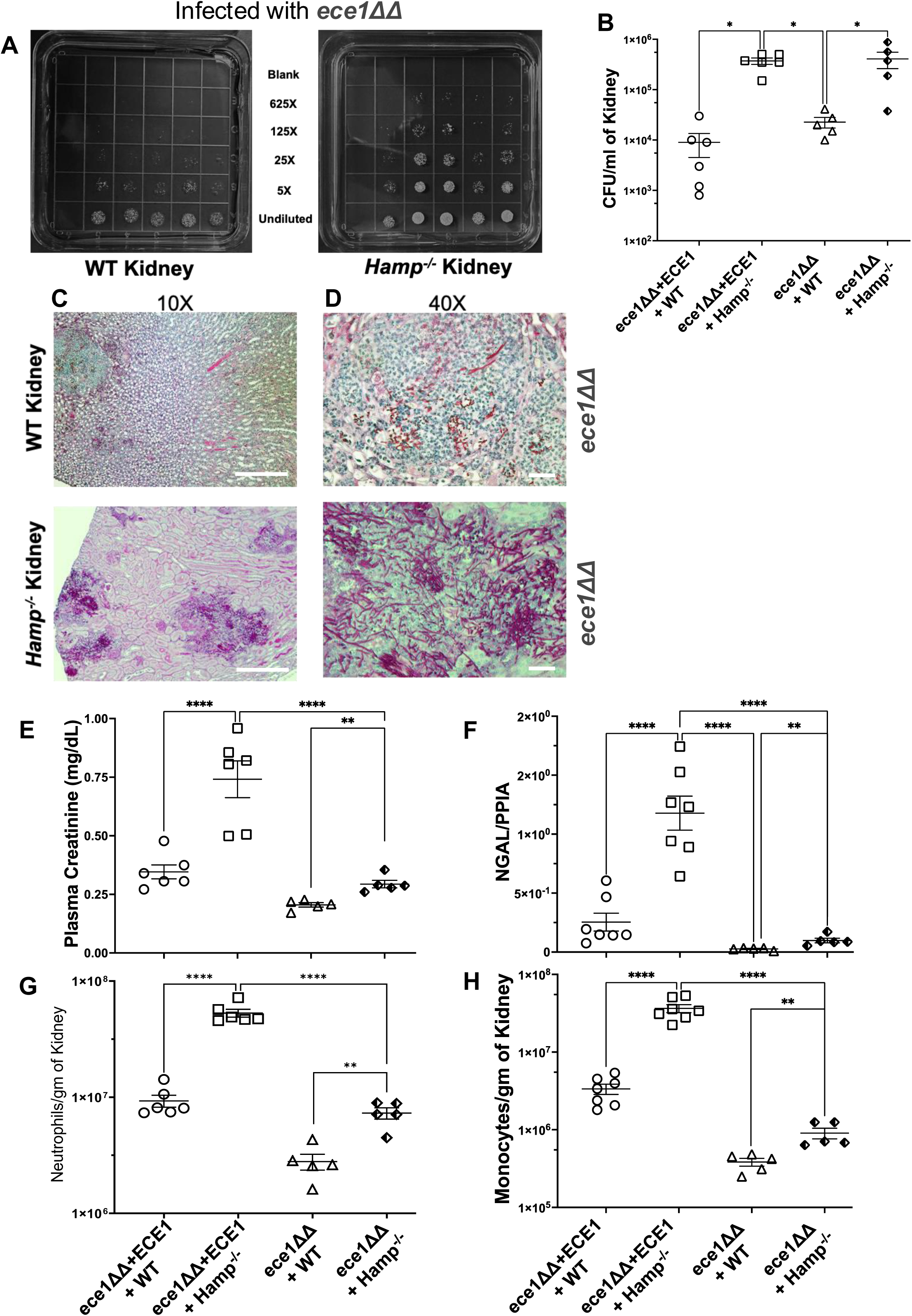
Hepcidin deficiency drives fungal burden, but the severity of AKI depends on candidalysin. W.T. and *Hamp^-/-^*mice were infected with 2e^5^ *Ece1* deficient *C. albicans* yeast (*ece1*ΔΔ), and the kidneys were harvested on day 3. Compared to W.T. kidneys, the fungal burden was significantly higher in the *Hamp^-/-^* kidneys (**A-B**) and was comparable to *Hamp^-/-^* kidneys infected with an isogenic strain sufficient for candidalysin (*ece1*ΔΔ *+ ECE1)* (**B**). The *ece1*ΔΔ *C. albicans* had undergone a hyphal transformation in *Hamp^-/-^* kidneys and histology showed multiple foci of infection, which were not observed in W.T. kidneys (**C-D**). Scale bar C, 10X:100 μM, D, 40X: 40 μM). Compared to W.T. mice, *ece1*ΔΔ strain infected *Hamp^-/-^* mice developed significantly more AKI as measured by plasma creatinine and intra renal *Ngal* gene expression (**E-F**). AKI was exacerbated by the *ece1*ΔΔ *+ ECE1* strain (**E-F**). Compared to W.T. kidneys, *ece1*ΔΔ infected *Hamp^-/-^* kidneys had significantly greater neutrophil and monocyte infiltrates (**G-H**). However, both the immune cell types were significantly lower than *ece1*ΔΔ *+ ECE1* infected *Hamp^-/-^* kidneys (**G-H**). Data was analyzed using 2-way ANOVA with Holm-Šídák’s multiple comparisons test and represented as mean ± SEM. *P < 0.05, **P < 0.005, ***P < 0.001. ****P < 0.0001.

### Iron exposes fungal **β**-glucan and promotes inflammation in the absence of candidalysin

To mechanistically identify how the *ece1Δ/Δ* strain was still able to induce kidney injury in *Hamp^-/-^*mice, we characterized the *ece1*ΔΔ*+ECE1* revertant and *ece1*Δ/Δ mutant cells grown under standard conditions or in the presence of high iron. Both strains grew comparably in the yeast morphology under standard conditions. However, the iron-rich broth sustained hyphal morphogenesis and exposed β-1, 3-glucan on the cell surface of both strains (**Fig. 6A-D**). Next, we evaluated the effect of iron-exposed *ece1*Δ/Δ yeast cells on HK-2 cells (a human renal PTEC cell line) under hyphal forming conditions (37^0^ C). Two hours after co-culture, the yeast cells transformed into hyphae, and the HK-2 cells secreted IL-8 in response to *ece1*Δ/Δ cultured under both conditions (**Fig. 6E**), indicating recognition of the fungus. However, fungal cells cultured under high iron-conditions induced significantly higher IL-8 secretion at both 2- and 4 hours post-infection as compared to cells grown under standard conditions (**Fig. 6E**). Thus, iron- induced changes in *C. albicans* elicit a greater inflammatory response from renal parenchymal cells independent of candidalysin. The *ece1*Δ/Δ subcultures grown in HK-2 medium with or without iron, under hyphal forming conditions (37^0^ C) grew comparably and ruled out increased fungal burden as a cause of higher IL-8 secretion (**Fig. 6F**). These *in vitro* studies shed light on the observed differences in renal pathology of W.T. and *Hamp^-/-^*mice infected with candidalysin- deficient *C. albicans* cells. These observations were validated *in vivo* using red fluorescent *C. albicans* (CAF2-1-dTomato)^69^ infected W.T. and *Hamp^-/-^* mice. Whole kidney homogenates of W.T. mice infected with the CAF2-1-dTomato strain revealed yeast and occasional red fluorescent hyphae with exposed β-1, 3-glucan (**Supplementary** Figure 6 **A, C, E**). In contrast, the infected kidney homogenates of *Hamp^-/-^*mice had significantly more yeast cells and hyphae showing higher levels of exposed β-1, 3-glucan (**Supplementary** Figure 6 **B, D, E**).

**Figure. 6:**
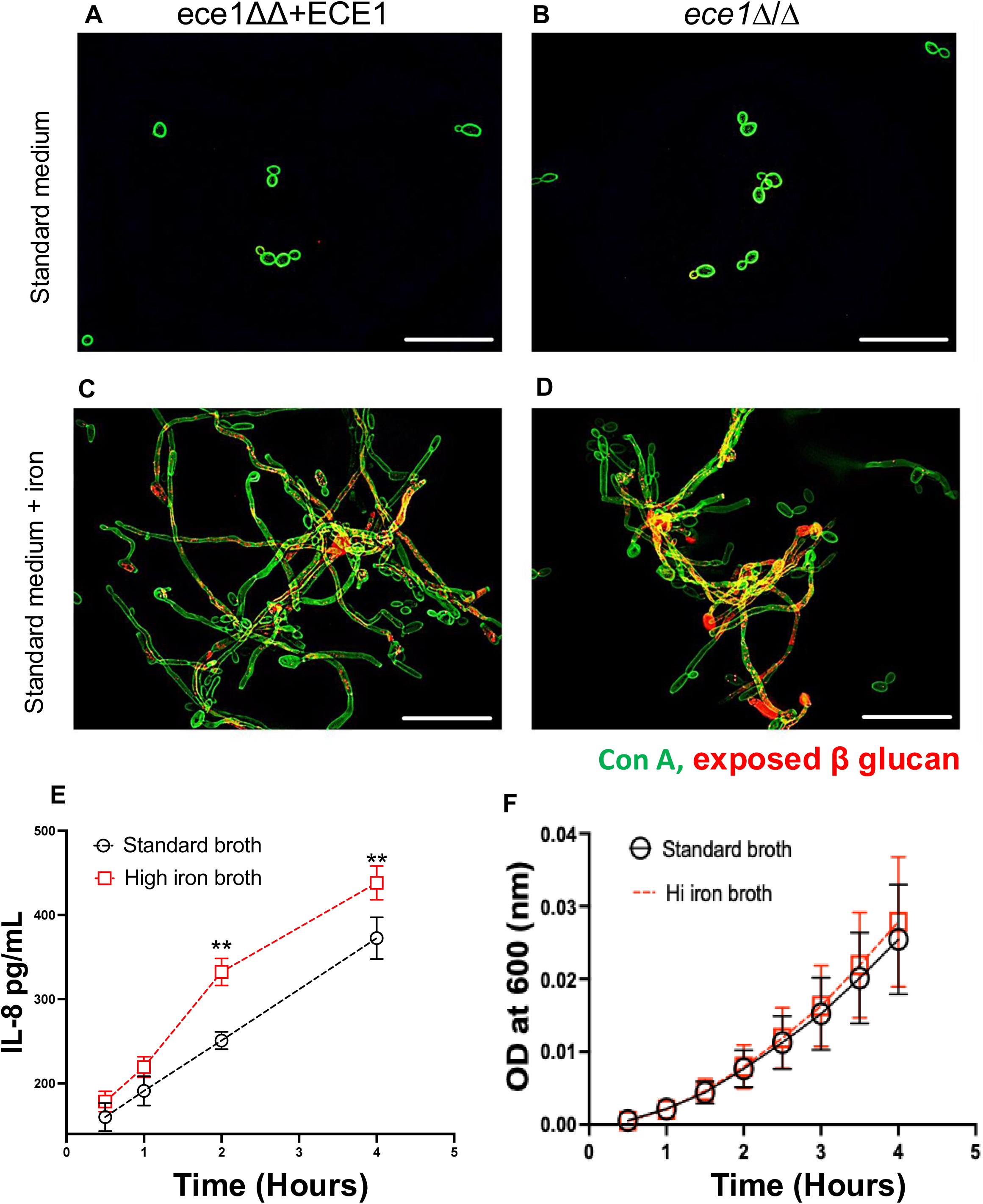
Iron exposes fungal **β**-glucan to potentiate chemokine production in human proximal renal tubular cells independent of candidalysin. *Ece1* knockout *C. albicans* yeast (*ece1*ΔΔ) and its isogenic candidalysin revertant (*ece1*ΔΔ *+ ECE1*) were grown in standard or iron-rich medium (100 μM FeCl_3_, equivalent to serum iron content of *Hamp^-/-^* mice) for twenty-four hours and stained for exposed β-glucan and Concanavalin A. Both fungal strains remained as yeast in the standard medium and did not expose β-glucan (**A-B**). However, adding iron to the growth medium transformed and sustained both *ece1*ΔΔ *and ece1*ΔΔ *+ ECE1* in their hyphal form with exposed β-glucans (**C-D**). Scale bar: 100 μM. The *ece1*ΔΔ *C. albicans* were grown overnight in standard or iron-rich medium (100 μM FeCl_3_) at 33^0^ C and then co-cultured with HK-2 cells at a MOI of 1:3 (Cells: Fungus) at 37^0^ C. Compared to *ece1*ΔΔ *C. albicans* grown in standard medium, ones grown in iron-rich medium induced significantly more IL-8 secretion (**E**). Data was normalized to IL-8 secreted by naïve cells at each time point. The growth of *ece1*ΔΔ *C. albicans* originally grown in standard and iron-rich medium and subsequently in HK-2 medium was followed at 37^0^ C for four hrs. There was no significant difference between the growth rate of the fungus in the two conditions (**F**). A 2-tailed Mann-Whitney test determined statistical significance at each time point and plotted as mean ± SEM. *P < 0.05, **P < 0.001.

### Acute hepcidin deficiency worsens *C. albicans*-induced pyelonephritis and can be rescued by synthetic hepcidin agonist

To evaluate the effect of acute, adult-onset hepcidin deficiency on outcomes of *C. albicans* infections, we used transgenic mice with tamoxifen-sensitive conditional *Hamp*1 deletion (termed iHamp*^-/-^*) mice^70^. These mice grow to adulthood with normal iron stores and serum iron parameters. However, upon exposure to Tamoxifen, *Hamp1* is irreversibly deleted, leading to iron overload in the serum, followed by the liver^70^. Compared to littermates, hepcidin production was suppressed following tamoxifen administration (**Fig. 7A-B**). As described previously, hepcidin deficiency was sufficient to deplete iron stores from the splenic red pulp (**Fig. 7C-D**). This study identifies that acute hepcidin deficiency is sufficient to initiate kidney iron loading (**Fig. 7E-F**). Following tamoxifen treatment and *C. albicans* infection, serum hepcidin increased in littermate controls (**Fig. 8A-B**). However, tamoxifen-induced deletion of the *Hamp* gene in i*Hamp^-/-^* mice failed to elicit the antifungal hepcidin response. Compared to littermates, the renal fungal burden on day four post-infection was significantly higher in i*Hamp^-/-^* mice (**Fig. 8C-D**), associated with hyphal formation in the kidneys (**Fig. 8E-F**) and increased tubular injury (**Fig 8H**). To evaluate the importance of early hepcidin activity for resistance to *C. albicans* infection, we injected *iHamp^-/-^* mice post-infection with mini-hepcidin (PR73: a synthetic hepcidin mimetic). Mini-hepcidin has greater potency and a longer-lasting effect than full-length hepcidin^71^. Daily administration of mini hepcidin significantly decreased the renal fungal burden (**Fig. 8C-D).** Histology revealed a few inflammatory foci and hyphae (**Fig. 8G, black dotted circle**) in mini hepcidin-treated mice. However, these were markedly less compared to vehicle controls. Less fungal burden was associated with attenuated PTEC injury as measured by intrarenal *Ngal* gene expression (**Fig. 8H**).

**Figure. 7.**
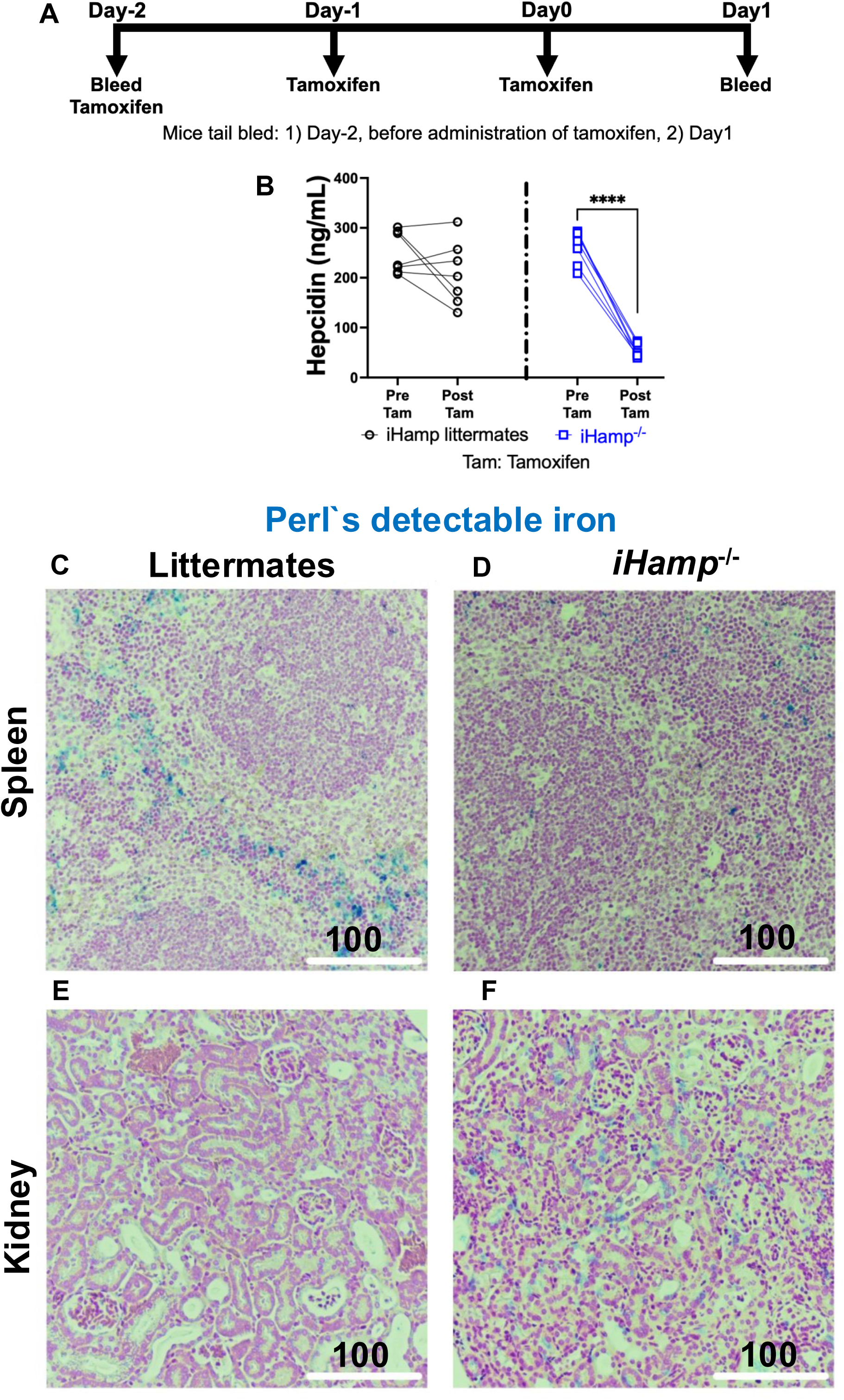
Acute hepcidin deficiency triggers iron loading in the kidneys. 10–12-week-old male and female inducible Hepcidin knockout mice (*iHamp^-/-^*) and their littermate controls (Cre -ve) were injected with 1mg Tamoxifen in corn oil for three consecutive days (**A**). The mice were tail-bled before the Tamoxifen injection. Serum hepcidin levels were measured 24 hr after the last dose of Tamoxifen. Before Tamoxifen injection, serum hepcidin levels in the *iHamp^-/-^* and their littermate controls were comparable but dropped significantly post-tamoxifen injection only in the *iHamp^-/-^*mice (**B**). Data was analyzed using paired T-test with Mann Whitney correction and are plotted as mean ± SEM. ****P < 0.0005. After tamoxifen injections, the formalin-fixed spleens and kidneys of littermate controls and *iHamp^-/-^*mice were stained for Perl’s detectable iron deposits. The littermate spleens stained positive for blue iron deposits in the red pulp region (**C**). However, the spleens of *iHamp^-/-^* mice lacked iron (**D**). The littermate kidneys did not show any Perl’s detectable iron (**E**), whereas the kidneys of *iHamp^-/-^* mice had started accumulating iron (**F**). Scale bar: 100 μM.

**Figure. 8:**
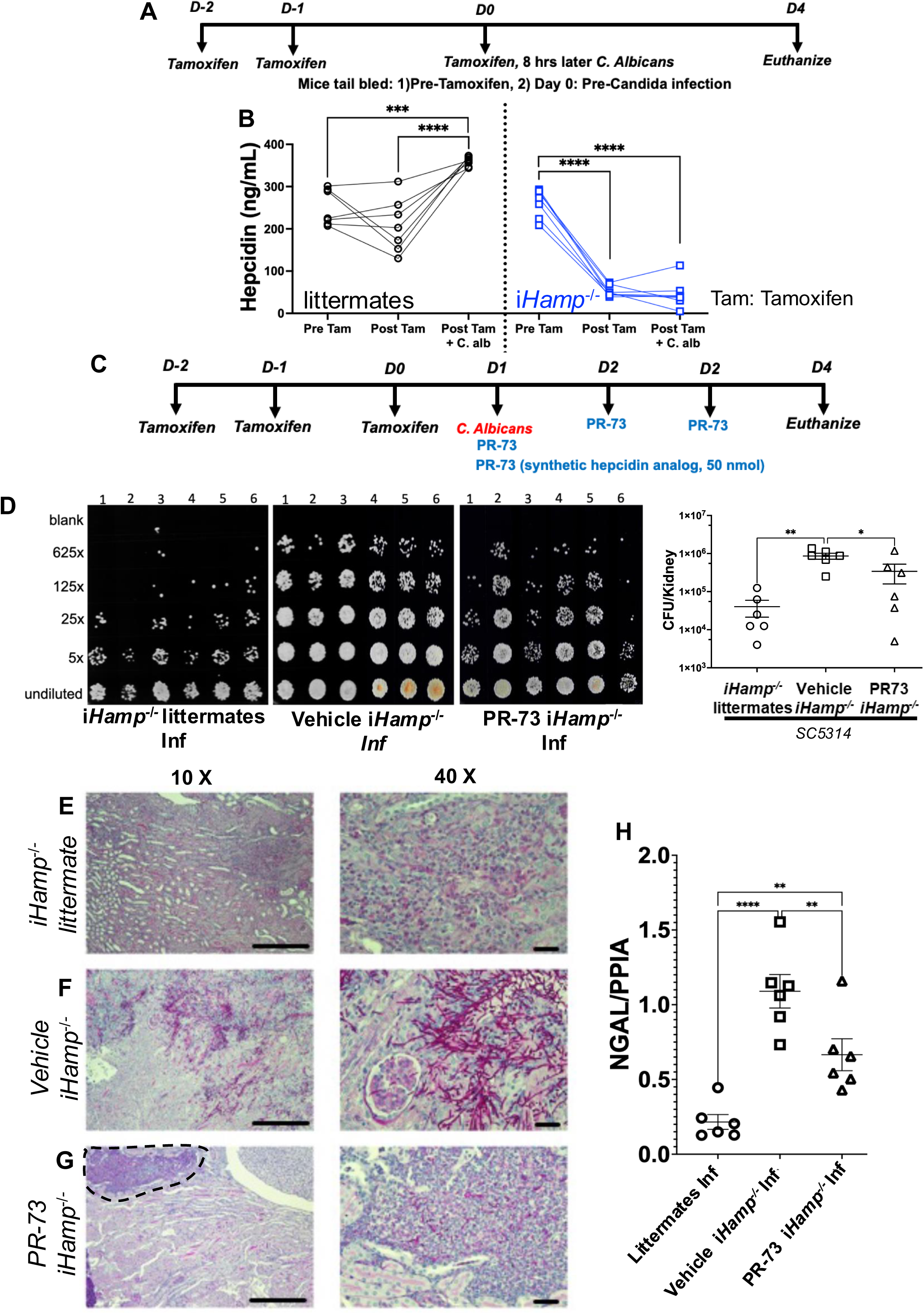
Acute hepcidin deficiency exacerbates *C. albicans*-induced renal pathology and is attenuated by PR-73, a synthetic hepcidin mimetic. Inducible hepcidin knockout mice (*iHamp^-/-^*) and their W.T. littermates were injected with Tamoxifen, as shown in **Fig. A**. Tamoxifen injection did not attenuate serum hepcidin levels in W.T. mice, which increased significantly post *C.* albicans infection (**B**). In contrast, Tamoxifen significantly attenuated serum hepcidin levels in *iHamp^-/-^* mice, which remained low after *C. albicans* infection (**B**). Statistical significance was determined by a 2-tailed Wilcoxon matched pair sign rank test and plotted as mean ± SEM. ***P < 0.001, ****P < 0.0001. W.T. littermates and *iHamp^-/-^* mice were injected with tamoxifen and *C. albicans*. The *iHamp^-/-^*mice received vehicle or PR-73 (50 nmol, intraperitoneal), starting 4 hours after candida infection (**C**). Compared to W.T. littermates, the kidneys of vehicle-treated *iHamp^-/-^* mice had a significantly higher fungal burden and was significantly reduced by PR-73 therapy (**D**). The fungus was still in the yeast form in the W.T. littermates (**E**) but had undergone a hyphal transformation in the vehicle-treated *iHamp^-/-^* mice (**F**). PR-73 treatment was associated with a reduction in hyphal morphogenesis (**G**). Reduced fungal burden and hyphal transformation in PR-73 treated *iHamp^- /-^*mice mitigated PTEC injury as measured by renal *Ngal* gene expression (**H**). Data was analyzed using 2-way ANOVA with Holm-Šídák’s multiple comparisons test and represented as mean ± SEM. *P < 0.05, **P < 0.005, ****P < 0.0001. Scale bar E-G: 100 uM and 30 uM.

## DISCUSSION

Herein, we identify an essential role of hepcidin in resistance to systemic *C. albicans* infections. We show that hepcidin deficiency and associated kidney iron assimilation worsen the outcome of disseminated candidiasis. Mechanistically, renal iron overload 1) promotes inexorable fungal growth, 2) sustains hyphal morphogenesis, and 3) exposes fungal wall components such as β-glucans to trigger inflammation and tissue injury independent of candidalysin in immunocompetent mice. Our conclusions are supported by genetically and inducible hepcidin-deficient mice, human observations, and *in vitro* studies corroborating a phenotype observed in chronic liver disease (CLD) patients. Of translational significance, we uncover hepcidin mimetic as an intervention and ferroportin as a therapeutic target to improve outcomes of disseminated candidiasis.

We show that the loss of hepcidin compromises nutritional immunity, instigating early uncontrolled fungal growth, filamentation, and mortality. Early *C. albicans* growth restriction by renal phagocytes protects against systemic candidiasis^69,72,73^. Hence, *C. albicans* initiates an iron acquisition program to survive in the hostile host environment^29,35,74^. Following exposure to *C. albicans,* intrarenal hepcidin levels increase^75^, and an iron import program is initiated in monocytes^76^ to limit access to iron, thereby mediating nutritional immunity. In support of this hypothesis, *C. albicans* isolated from W.T. murine kidneys have an iron starvation profile^77^. However, the loss of hepcidin leads to iron overload of the renal epithelium and negates the iron retention ability of monocytes and neutrophils^78–80^, providing unhindered access to this key nutrient. We also show that hepcidin deficiency is associated with a hyper-inflammatory phenotype in the *C. albicans*-infected kidneys. Work from our lab has demonstrated that hepcidin attenuates LPS and TLR3-induced immune responses in mice and macrophages^81,82^. Our study sets the stage for defining the functional consequences of monocyte and neutrophil- specific intracellular iron deficiency or excess during *Candida* infections.

Prior work has shown that candidalysin is essential for epithelial damage^61,83,84^, inflammatory response^85^, and renal neutrophil recruitment^68^ during *C. albicans* infections *in vitro* and *in vivo*. We uncover a previously unknown hepcidin-iron-dependent mechanism that promotes fungal virulence and induces renal injury independent of candidalysin in immunocompetent mice. We demonstrate that iron sustains *C. albicans* hyphae with exposed β- 1, 3-glucan and accentuates IL-8 production in human renal epithelial cells. Exposed β-glucans on *C. albicans* cell surfaces are recognized by oral epithelial cells via the ephrin type-A receptor 2 (EphA2) ^86^ and trigger an inflammatory response. EphA2 is also expressed by renal tubular epithelial cells^87–89^ and can account for candidalysin-independent renal pathology observed in *Hamp^-/-^* mice. Furthermore, neutrophil EphA2 also serves as a receptor for β-glucans^90^ and may explain the high neutrophil numbers in *ece1*Δ/Δ infected *Hamp^-/-^* mice. Dectin-1 also recognizes exposed β-1, 3-glucan and is widely expressed on cells of the myeloid lineage and different types of epithelial cells ^91^. However, its expression on the renal epithelium has not been reported, and the kidney resident macrophages and dendritic cells may initially play out this role. We acknowledge that excess iron can trigger additional changes to potentiate virulence in *C. albicans* not identified in this study. Isolating and profiling *C. albicans* from *Hamp^-/-^* mice can provide clues and merits investigation in future studies.

A key aspect of our research is its extension to human disease, as indicated by (1) attenuated hepcidin production in humans and mice with liver fibrosis; (2) identifying that liver fibrosis is associated with renal iron accumulation, (3) the ability of PR73, a hepcidin mimetic^92,93^ to ameliorate outcomes of invasive *C. albicans* infection. We have previously shown that hepcidin deficiency does not influence the outcome of aspergillosis^94^ and as such our findings appear relevant to specific human infections and disease states. For example, patients with chronic liver diseases are at a higher risk of candidemia than patients without liver disease, even in the absence of neutropenia^51–53^. However, the underlying mechanisms are incompletely defined. Our findings suggest that iron may accumulate in the tissues of CLD patients due to low hepcidin levels and render them susceptible to fungal infections, summarized in a schematic model (**Figure 9**). While we could not obtain kidney biopsies of CLD patients, the findings in *iHamp^-/-^* mice and the conserved role of hepcidin in humans and mice provide a strong rationale. Our findings are also relevant to patients harboring loss-of-function *STAT3* variants (Job’s syndrome), where care is directed at treating and preventing recurrent fungal infections^95^. Phosphorylation of STAT3 is critical for hepcidin transcription^47,49,50^, and the link between *STAT3* variants, hepcidin, and recurrent fungal infections has not been explored.

**Figure. 9:**
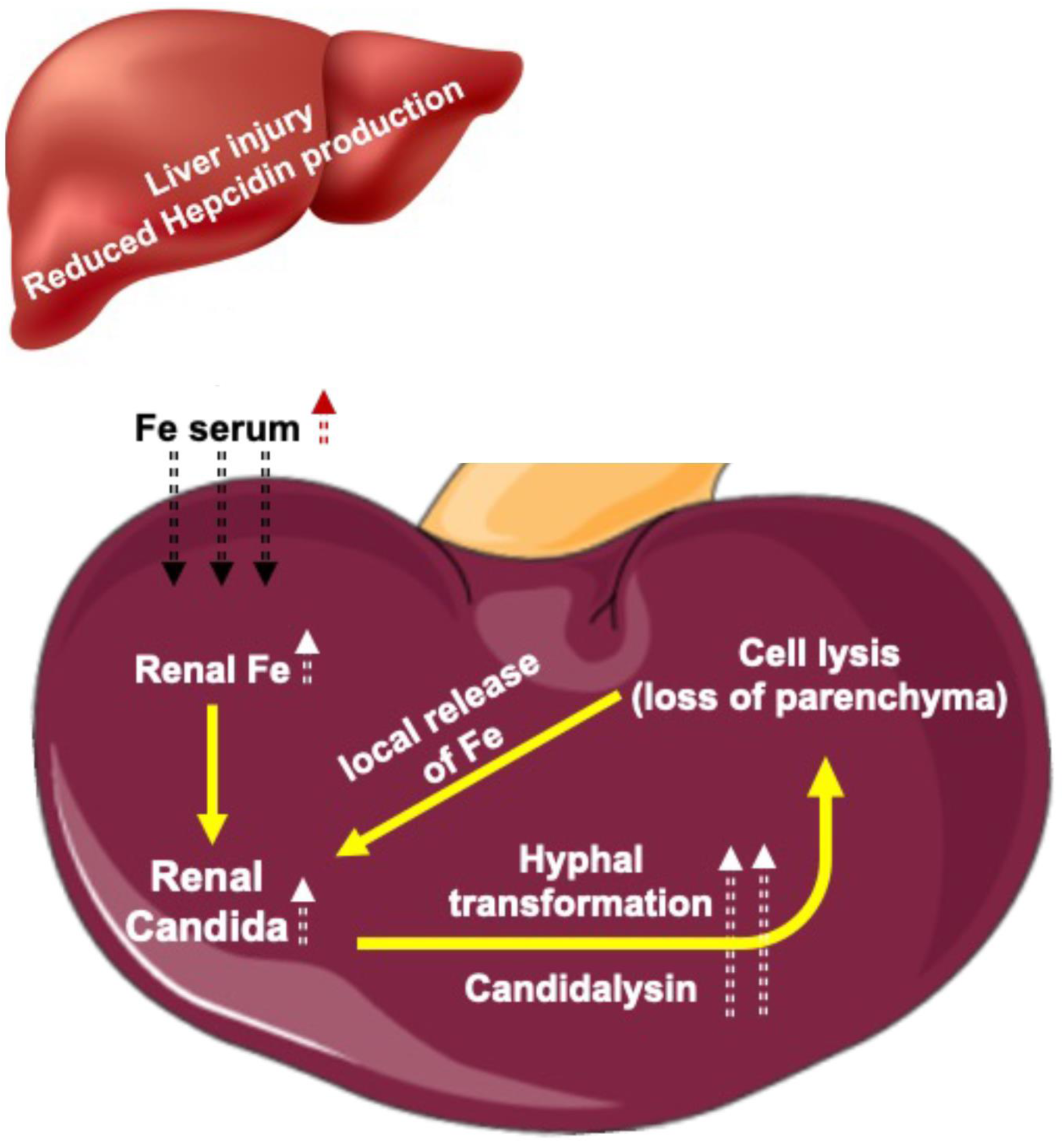
**Proposed model**: We propose that chronic liver disease attenuates hepcidin production, impairs organismal control of iron regulation, and increases iron accumulation in organs including the kidneys. *C. albicans* is trophic to the kidneys and undergoes inexorable growth and virulent hyphal transformation in an iron-rich environment. The candidalysin- secreting hyphae lyse the iron-loaded parenchymal cells, promote a local microenvironment that propagates fungal growth, and perpetuate kidney injury.

In summary, our study provides mechanistic and translational insights into the essential role of hepcidin in orchestrating nutritional immunity during fungal disease, thereby uncovering hepcidin as a novel host susceptibility factor in resistance to systemic candidiasis.

## TRANSLATIONAL STATEMENT

Hepcidin is a promising intervention for developing individualized risk stratification and prognostication strategies. As an endogenous peptide with a favorable safety profile (ClinicalTrials.gov identifier: NCT03381833 and NCT03165864), hepcidin and its analog carry a high translational potential. In the era of increasing fungal infections that continue to exhibit poor outcomes despite conventional antifungal therapies (https://fis.fda.gov/sense/), our studies lay the foundation to evaluate novel hepcidin mimetics such as Vamifeport^96,97^ or Rusfertide^98,99^, as adjunct therapies to mitigate fungal infections and improve the prognosis of at-risk patients.

## ACKNOWLEDGMENT

This work was supported by the NIH (grant # RO1DK136011) and Vifor Pharma (grant # P0213104) to Y.S. and NIH (grant # RO1AI135128) and Keck Foundation (grant # 994413) to B.M. This work was supported in part by the Division of Intramural Research of the NIAID, NIH (MSL).

## AUTHOR CONTRIBUTION

YS conceived the study, and Y.S., B.M., and MSL designed the experiments. T.A., D.K., SK, D.D., TA, N.K., B.M., and Y.S. performed the experiments. S.V. generated and provided the Hepcidin knockout mice. B.H. provided the *ece1*ΔΔ*+ECE1* and *ece1*1/1 *C. albicans* strains and gave expert advice. EN provided PR-73 (mini hepcidin) and expertise on its usage. A.D. provided the inducible hepcidin knockout mice and advice on their phenotyping. MSL provided experimental protocols, discussed the results, and gave expert advice. T.A., D.K., and Y.S. analyzed the data. T.A. and Y.S. wrote the original draft of the manuscript. All authors edited the manuscript and approved the final version.

## Supplementary Figure

**Supplementary** Figure 1: Hepcidin deficiency depletes splenic iron stores and is associated with kidney iron overload.

**Supplementary** Figure 2: Characterization of W.T. and *Hamp^-/-^* tissue post *C. albicans* infection

**Supplementary** Figure 3: Renal iron content accelerates fungal growth and sustains its hyphal state.

**Supplementary** Figure 4. *C. albicans* infection in *Hamp^-/-^* mice is associated with increased intra-renal gene expression of inflammatory cytokines and chemoattractants.

**Supplementary** Figure 5. Gating strategy to identify intra-renal neutrophils and monocytes.

**Supplementary** Figure 6. In vivo detection and quantification of exposed β-1, 3-glucan.

## STAR **IZ** METHODS

Detailed methods are provided include the following:

- KEY RESOURCES TABLE
- RESOURCE AVAILABILITY

`o` Materials availability
- EXPERIMENTAL MODEL AND SUBJECT DETAILS

`o` Human subjects
`o` Mice
`o` Fungal strain and mouse model of systemic candidiasis
- METHOD DETAILS

`o` Induced liver fibrosis model
`o` Fungal burden determination
`o` Biochemical assays and tissue samples
`o` PAS and H&E analysis
`o` Grocott Methenamine silver
`o` Perls staining
`o` Plasma creatinine assay
`o` Immunofluorescence
`o` In vitro detection of exposed β-1,3-glucan
`o` In vivo detection of exposed β-1,3-glucan
`o` Flow cytometry
`o` Real-time PCR
`o` Influence of iron on the growth of *C. albicans* in mouse kidneys and human PTEC cell lysates
`o` Effect of iron on *C. albicans* hyphal sustenance
`o` Effect of *ece1*ΔΔ *C. albicans* grown in excess iron on human proximal renal tubular cells immune response

