## Supplemental Figures for "Essential role of Hepcidin in host resistance to disseminated candidiasis"

Supplemental Figure. 1

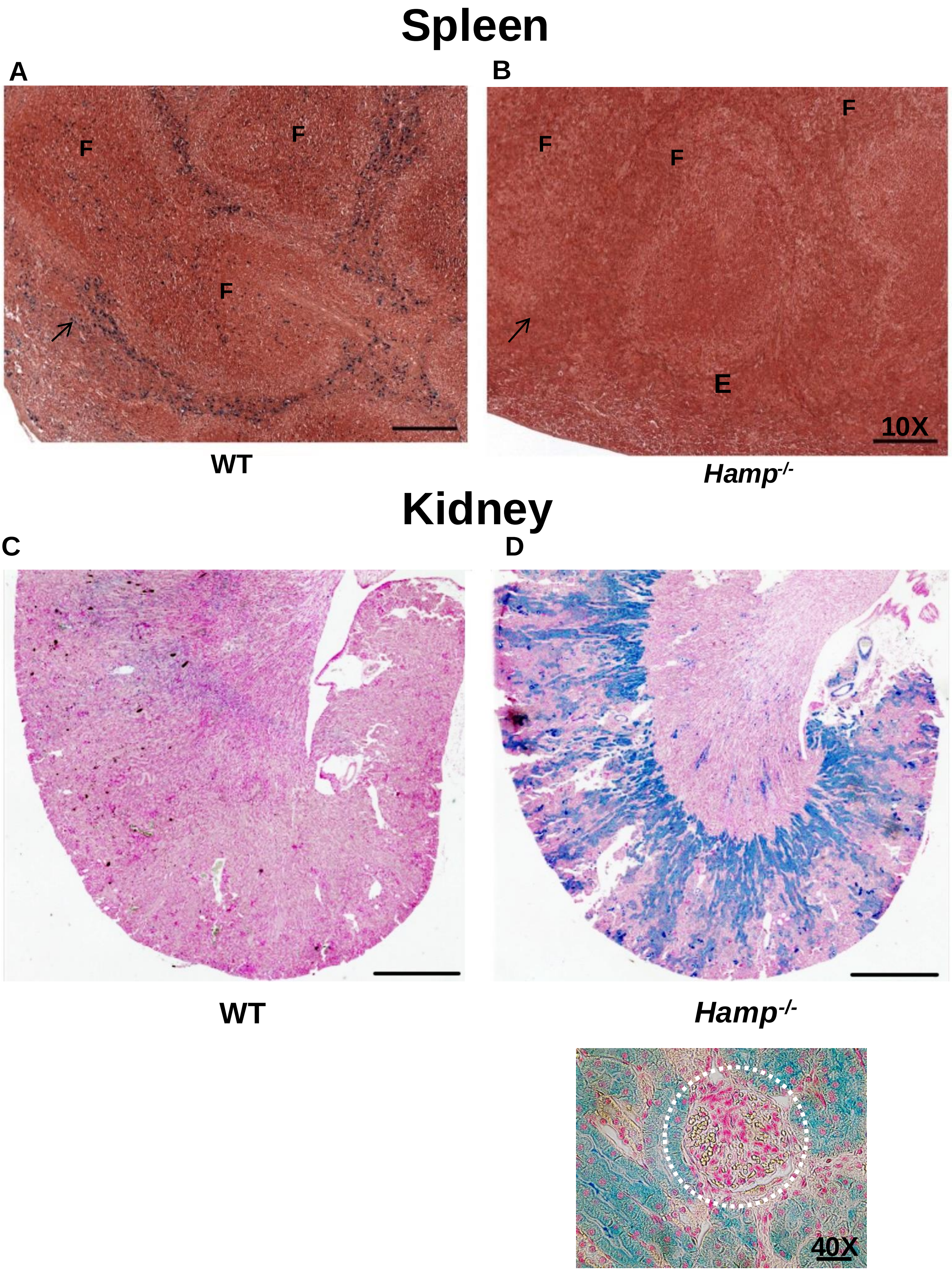

**S1: Hepcidin deficiency depletes splenic iron stores and is associated with kidney iron overload.**

Formalin-fixed spleens and kidneys of naive 10–12-week-old littermate controls (WT) and hepcidin knockout (*Hamp*$^{-/-}$) mice (all on B6 background) were stained for Perl's detectable iron deposits. The WT spleens stained positive for blue iron deposits in the red pulp region (**A**). However, the spleens of *Hamp*$^{-/-}$ mice lacked iron in the red pulp region (**B**). F: splenic follicle. In contrast, WT kidneys did not show any Perl's detectable iron (**C**), whereas the *Hamp*$^{-/-}$ kidneys had significant iron deposits that extended from the cortico-medullary region to the deep cortex (**D**). The glomeruli (within the white dotted line) were devoid of any observable iron deposits, and most of the iron deposits were in the tubular segments (**E**). Scale bar: 10X:100 μM, 4X:100 μM, and 40X: 30 μM.

Supplemental Figure. 2

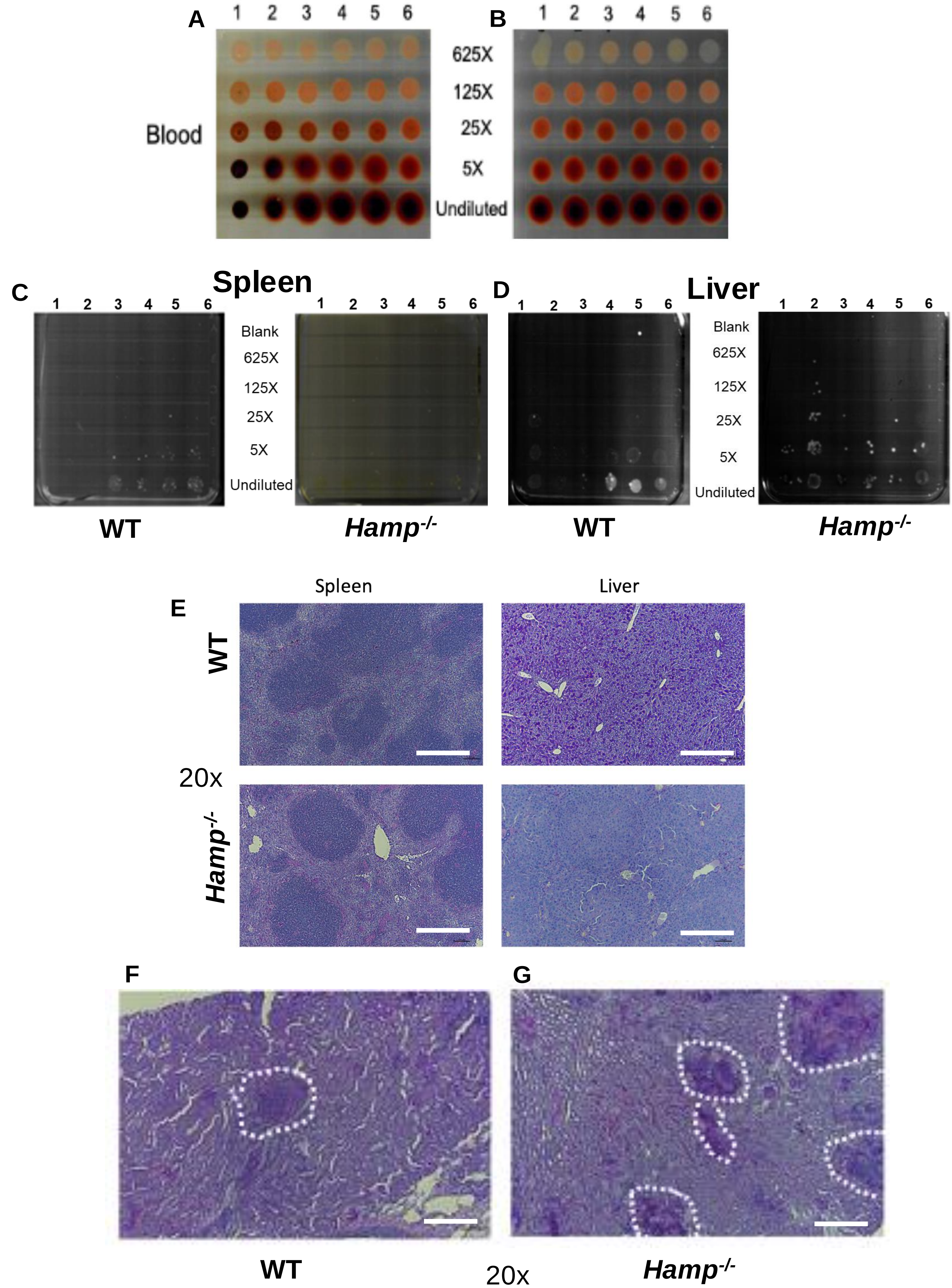

**S2: Characterization of WT and *Hamp*$^{-/-}$ tissue post *C. albicans* infection**

12-week-old WT and *Hamp*$^{-/-}$ mice were intravenously infected with 2 X $10^5$ SC5314 yeast cells and tail bled after 24 hrs. At this time point, candidemia was not observed in either strain (**A-B**). 3-days post-infection fungal burden in iron-sufficient WT spleens was significantly higher than iron-depleted spleens of *Hamp*$^{-/-}$ mice (**C**). In contrast, the fungal burden in the iron-loaded livers of *Hamp*$^{-/-}$ mice was significantly higher (**D**) (Data is quantified in **Fig. 1G-H**). The PAS-stained spleen and liver sections revealed no obvious fungal growth or pathology (**E**). However, while the kidneys of WT mice displayed a few small granulomatous foci of inflammation with fungal abscesses (**F**), the *Hamp*$^{-/-}$ kidneys had multiple large granulomatous foci of inflammation with fungal abscesses (**G**). Scale bar: 50 μM.

#### Supplemental Figure 3

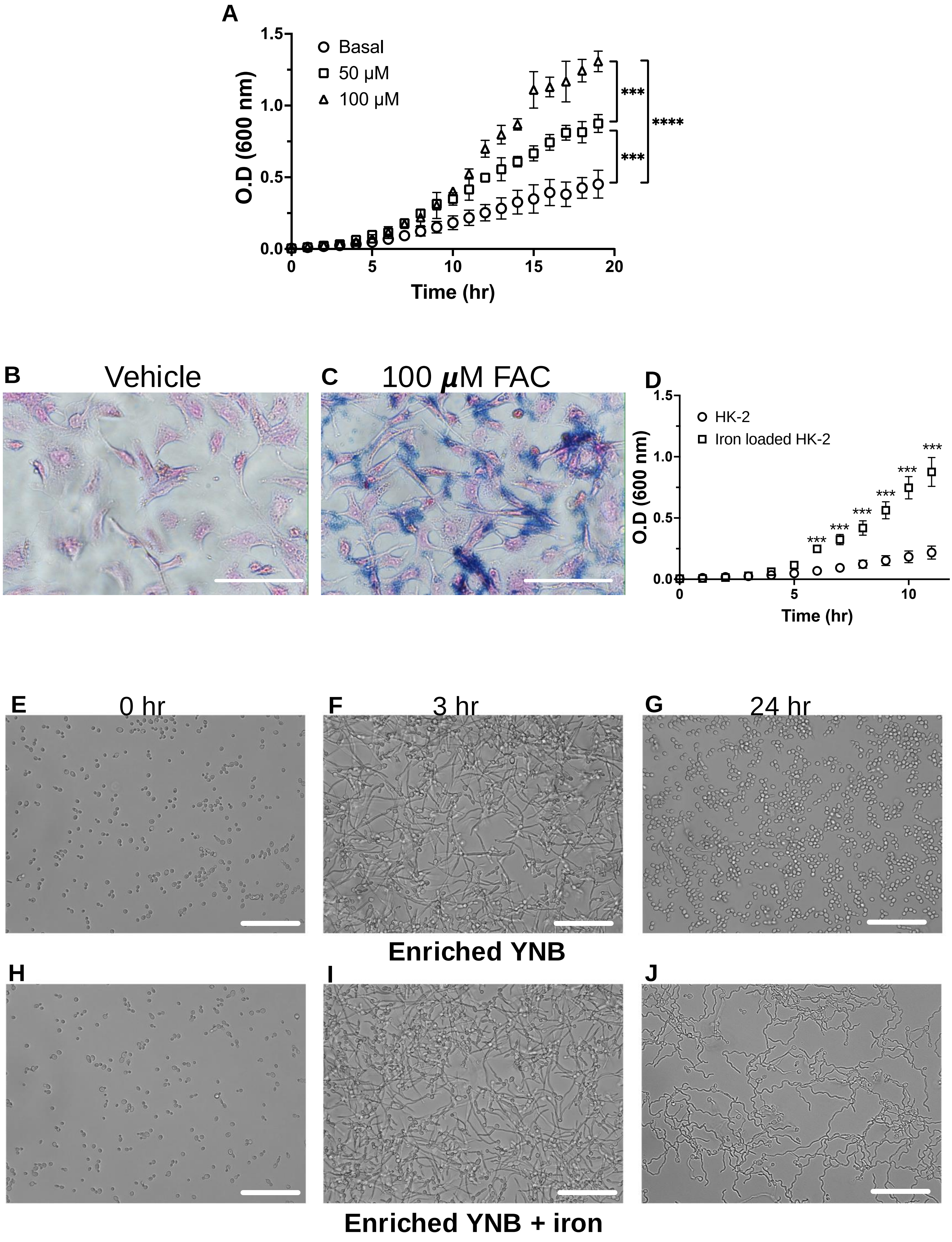

**S3: Renal iron content accelerates fungal growth and sustains its hyphal state.**

Single-cell suspension of perfused and red blood cell lysed WT kidneys were spiked with 50 or 100 μM ferric ammonium citrate (see Star Protocol for experimental details). The cultures were washed to remove cell-free iron lysed, and the supernatant was spiked with 10,000 *C. albicans* yeast cells. The growth curves were generated for 20 hours at $32^0$C. *C. albicans* growth increased with tissue iron content (**A**). HK-2 cells (human proximal tubular epithelial cell line) were vehicle or iron-loaded (100 ***μ***M ferric ammonium citrate (FAC)) for 48 hours and stained for iron deposits by Perl's blue staining **(B-C**). Vehicle or iron-loaded HK-2 cells were cultured with *C. albicans* for 12 hours at $37^0$C (MOI 1:1, cell: fungus). The supernatants were collected and spiked with an equal number of *C. albicans* cells (in yeast form). There was a significantly higher fungal growth in the supernatant of iron-loaded HK-2 cells (**D***). C. albicans* was grown overnight in YNB or YNB with 100 μM FAC at $33^0$C. After washing with fresh medium, one million yeasts from each condition were sub-cultured into 12-well plates, and their growth at $37^0$C was monitored at different time points. Yeast in both conditions had transformed into hyphae after three hours (**E-F, H-I**). However, after twenty-four hours, only the high iron broth sustained *C. albicans* in hyphal form (**G and J**). Images were taken at 40X magnification using Keyence BZ-X800 imaging microscope. Scale bar: 50 microns.

**Supplemental Figure 4**

### Inflammation

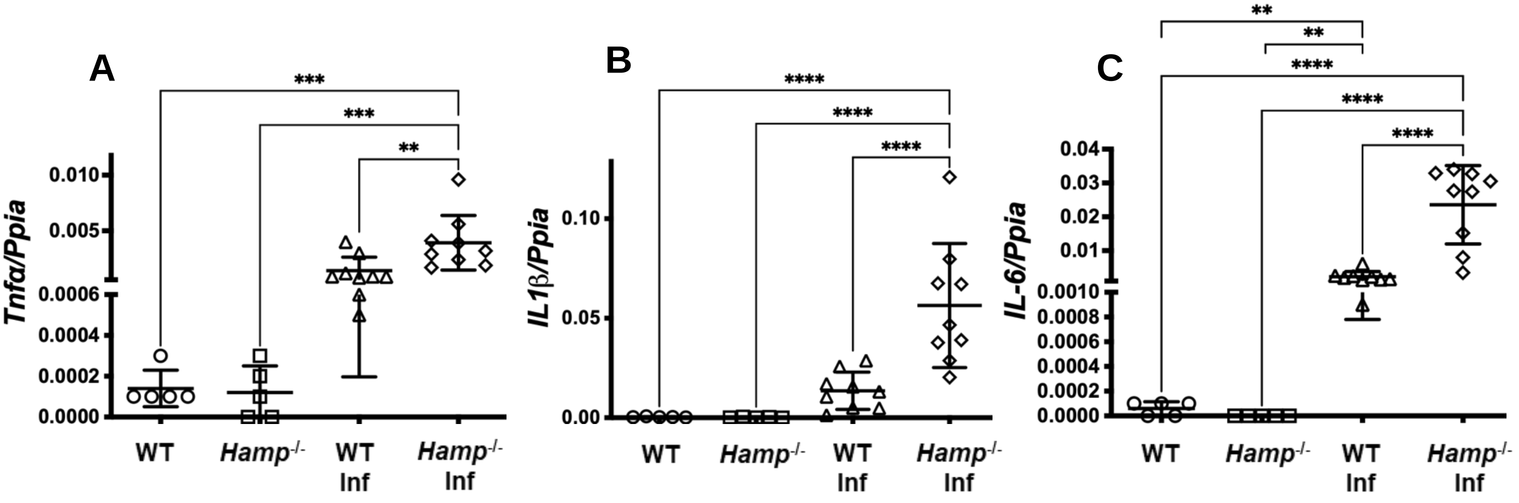


### Chemoattractant

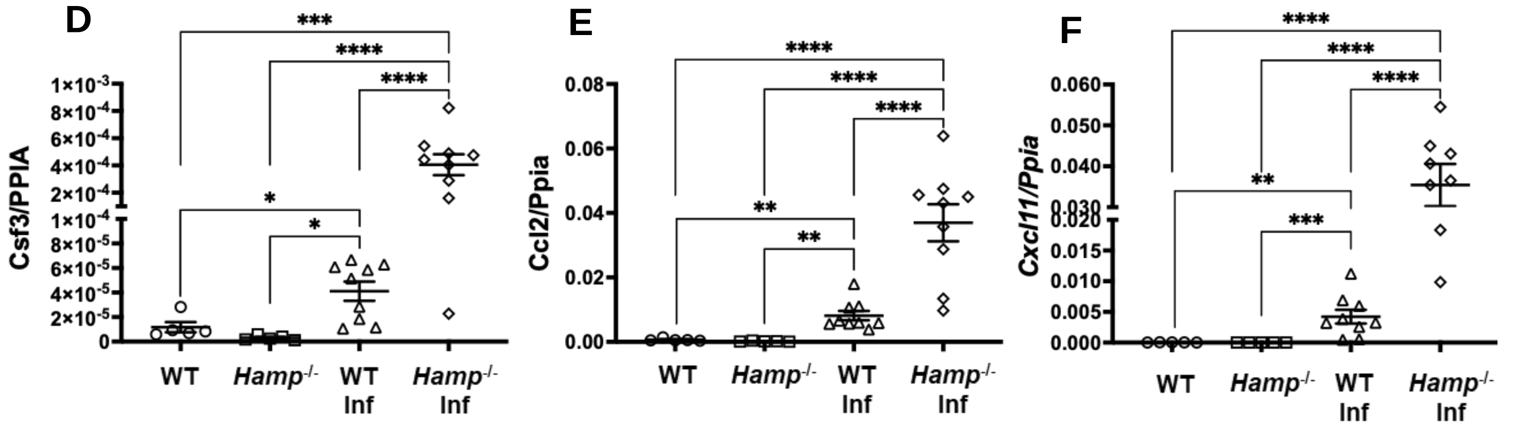

**S4. *C. albicans* infection in *Hamp*$^{-/-}$ mice is associated with increased intra renal gene expression of inflammatory cytokines and chemoattractants.**

WT and *Hamp*$^{-/-}$ mice were infected with 2e$^{5}$ *C. albicans* yeast, and tissues were harvested three days later. Naïve WT and *Hamp*$^{-/-}$ mice had comparable intra renal gene expression of *Tnf*$\alpha$ (**A**) *IL-1*$\beta$ (**B**) and *IL-6* (**C**) (inflammatory cytokines) and chemoattractants such as *Csf3* (**D**), *Ccl2* (**E**), and *Cxcl11* (**F**). However, compared to WT kidneys, *C. albicans* infection significantly increased the intra renal gene expression of inflammatory cytokines (**A-C**) and chemoattractant (**D-F**) in *Hamp*$^{-/-}$ kidneys. Data was analyzed using 2-way ANOVA with Holm-Šídák's multiple comparisons test and represented as mean ± SEM. *P < 0.05, **P < 0.005, ***P < 0.001. ****P < 0.0001.

#### Supplement Figure 5

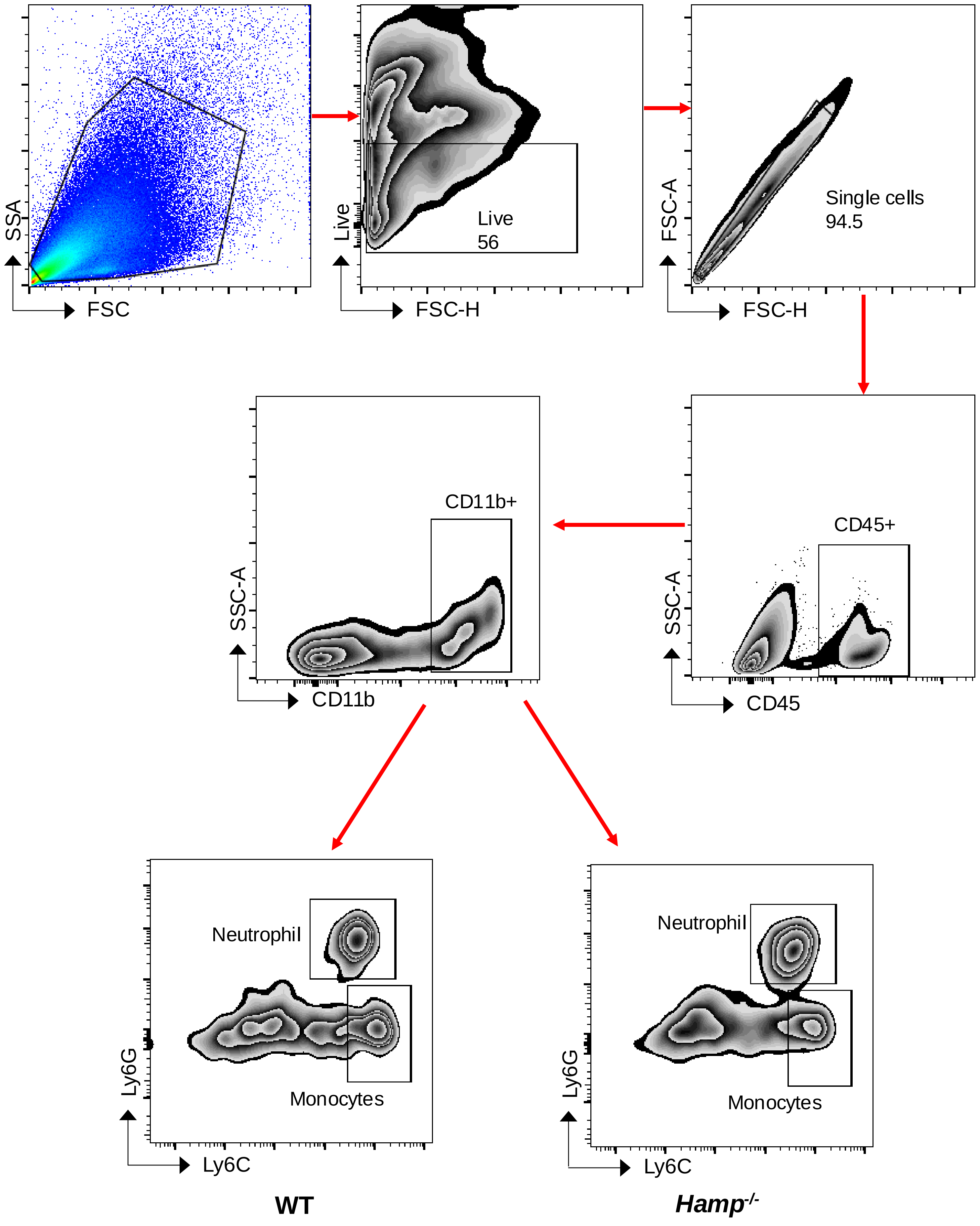

**S5. Gating strategy to identify intra-renal neutrophils and monocytes.**

Kidneys were stained with a fixable live/dead dye. After blocking FC receptors, antibodies were added to CD45, CD11b, Ly6C, and Ly6G (1A8). Neutrophils were identified as: Within live gate, singlets, CD45+ve, CD11b+ve, Ly6G+ve(1A8), and Ly6C+ve. Monocytes were identified as: Within live gate, singlets, CD45+ve, CD11b+ve, Ly6C+ve, and Ly6G-ve(1A8).

**Supplement Figure 6**

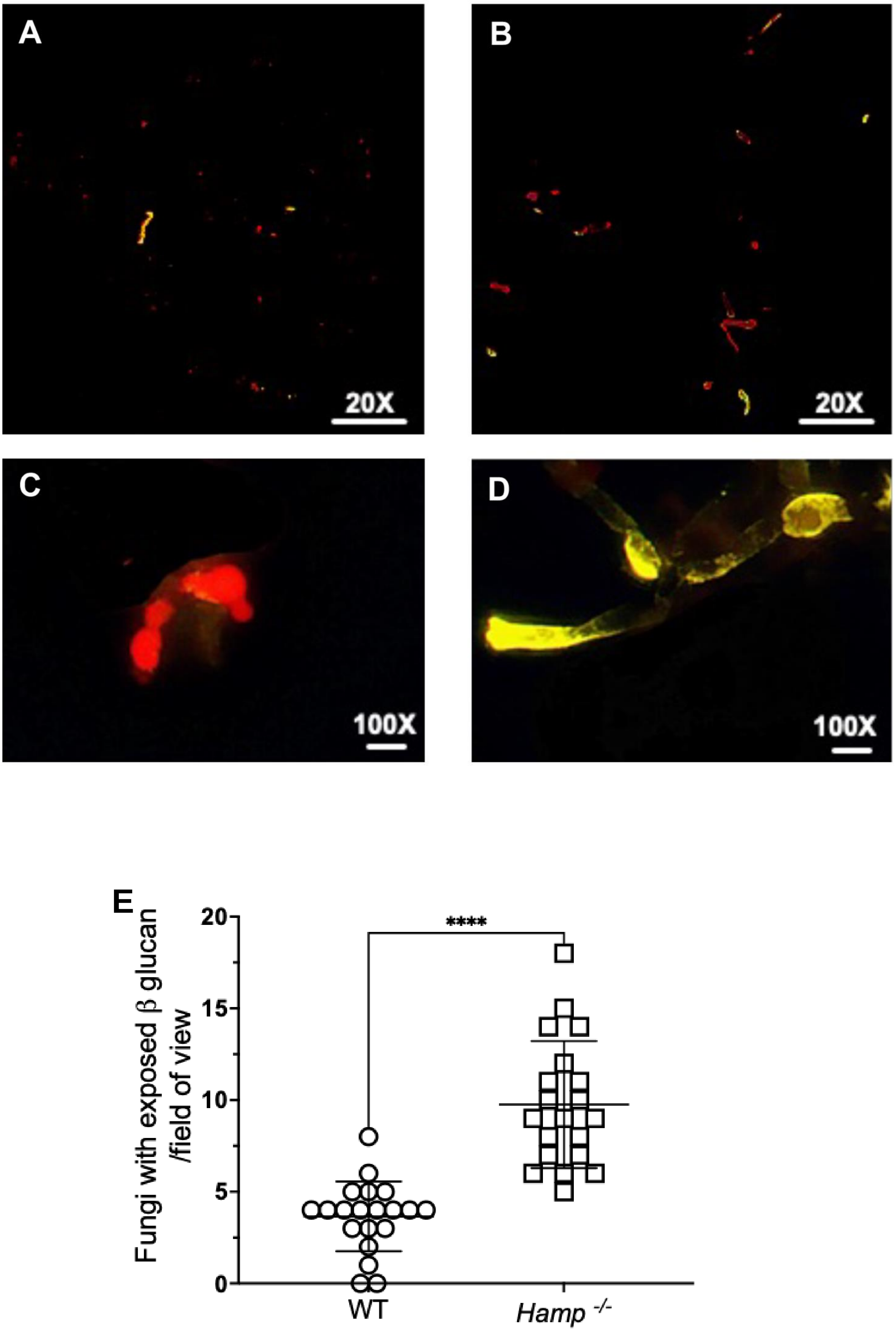

**S6. In vivo detection and quantification of exposed β-1, 3-glucan.**

WT and *Hamp*$^{-/-}$ mice were intravenously infected with two hundred thousand red fluorescent *Candida albicans* (CAF2-1-dTomato)(*69*). After three days, whole kidney homogenates were analyzed for fungal cells with exposed β-1, 3-glucan. Infected WT kidneys revealed red yeast and occasional hyphae with exposed β-1, 3-glucan (A and C). In contrast, the infected kidney digests of *Hamp*$^{-/-}$ mice had more red-fluorescent hyphae with exposed β-1, 3-glucan (green) (B and D). Scale bar A and B: 20X: 100 μM, C, and D: 100X: 30 μM. For quantifying fungi with exposed β-1, 3-glucan, five random 20X images from each slide (n = 5 from each strain) were captured. Each dot represents the number of yellow fungi within an image. Individual values are plotted with standard deviation (E). A 2-tailed Mann-Whitney test was used to determine statistical significance. ***$P < 0.0001$.
