## Supplementary Methods for "Essential role of Hepcidin in host resistance to disseminated candidiasis"

KEY RESOURCES TABLE

| REAGENT or RESOURCES | SOURCE | IDENTIFIER |
| --- | --- | --- |
| Chemicals |  |  |
| Yeast Peptone Dextrose Broth | Difco | DF0428-17-5 |
| Yeast Peptone Dextrose Agar | Difco | DF0427-17-6 |
| YNB Broth w/o ammonium sulfate, w/o copper sulfate w/o ferric chloride | MP Biomedicals | 4027112 |
| Copper sulfate, Pentahydrate | LabChem | LC134051 |
| Ferric Ammonium Citrate | Sigma | F5879 |
| Ammonium sulfate | Fisher chemical | A702-500 |
| Glucose | Sigma | G7021-100G |
| CSM (Powder) | MP Biomedicals | 4500012 |
| Penicillin/Streptomycin | Gibco | 15140122 |
| Ketamine | UF ACS |  |
| Xylazine | UF ACS |  |
| 10% Formalin | Fisher Brand | 245-684 |
| Tamoxifen | Sigma | T5648 |
| Triton-X-100 | Sigma Aldrich | 50-178-1841 |
| Trizol | Ambion | 15596018 |
| Chloroform | Sigma-Aldrich | 319988 |
| CCl_4_ Carbon Tetrachloride | Sigma | 289116 |
| Keratinocyte serum-free medium | Gibco | 17005042 |
| Bovine pituitary extract | Gibco | 13028-014 |
| Human recombinant epidermal growth factor | Gibco | 10450-013 |
| Renal epithelial cell growth basal medium 2 | PromoCell | C-26235 |
| **Supplement Pack** Renal Epithelial Cell GM2 | PromoCell | C-39605 |
| PR-73 mini hepcidin | Kind gift from Elizabeta Nemeth, UCLA, USA |  |
| ProLong Gold antifade agent with DAPI | Invitrogen | P36962 |
| ProLong Gold antifade agent without DAPI | Invitrogen | P36961 |
| Collagenase (Type IV) | Worthington | LS004188 |
| DNAse | Roshe | 04536282001 |
| Nuclear fast red |  |  |
| Commercial Assays |  |  |
| Creatinine Assay (Enzymatic) | Diazyme | DZ072B |
| Urea Nitrogen (BUN) Colorimetric | Arbor Assays | K024-H |
| RNeasy Plus Mini Kit | Qiagen | 74104 |
| IL-8 Human ELISA Kit | Invitrogen | KHC0082 |
| Grocott Methenamine Silver Stain (GMS) for Fungus & PCP | Polysciences | 25087-1 |

| Primers- Mouse | SOURCE | IDENTIFIER |
| --- | --- | --- |
| PPIA | Bio-Rad | qMmuCED0041303 |
| Tnfα | Bio-Rad | qMmuCED0004141 |
| IL-1β | Bio-Rad | qMmuCID0005641 |
| IL-6 | Bio-Rad | qMmuCEDD0045760 |
| GSDMD | Bio-Rad | qMmuCED0003802 |
| MLKL | Bio-Rad | qMmuCED0044462 |
| CSF3 | Bio-Rad | qMmuCED0004279 |
| CCL2 | Bio-Rad | qMmuCED0003785 |
| CXCl11 | Ori-gene | MP202411 |
| Fungal Primer Sequence |  |  |
| ECE1 forward | Integrated DNA technologies | atcgaaaatgccaagagag |
| ECE1 Reverse | Integrated DNA technologies | agcattttcaataccgacag |
| TDH3 Forward | Integrated DNA technologies | atcccacaaggactggaga |
| TDH3 Reverse | Integrated DNA technologies | gcagaagctttagcaacgtg |

| Antibodies |  |  |
| --- | --- | --- |
| PE/Cyanine7 anti-mouse CD45 (Clone 30-F11) | Biolegend | 103114 |
| FITC anti-mouse/human CD11b (Clone M1/70) | Biolegend | 101206 |
| PE anti-mouse Ly-6C (Clone HK 1.4) | Biolegend | 128008 |
| APC anti-mouse Ly-6G (Clone 1A8) | Biolegend | 127614 |
| (1-3)-beta-glucan-directed monoclonal antibody | Biosupplies Australia PTY LTD | 400-2 |
| Concanavalin A | Invitrogen | C11252 |
| Goat Anti-Mouse IgG, Human ads- PE | Southern Biotech | 1030-09S |
| Goat Anti-Mouse IgG- AF647 | Abcam | ab150115 |
| Fungal Strains | Source |  |
| *Candida albicans SC5314* | ATCC | MYA-2876 |
| *Candida albicans ece1ΔΔ*  *Candida albicans ECE1* revertant *ece1ΔΔ* **+** ECE1 | Bernhard Hube  (Leibniz Institute for Natural Product Research and Infection Biology – Hans Knöll Institute, Germany) |  |
| *Candida albicans CAF2-1-dTomato* | Michail Lionakis  (NIH, Bethesda)  Lionakis et. al., 2013^1^ | YCAT1127 |
| Experimental Model: Mouse strains | SOURCE |  |
| Hepcidin KO mice (Hamp^-/-^) | Sophie Vaulont  (Institut Cochin, France) |  |
| Inducible hepcidin KO mice  (iHamp^-/-^) | Alexander Drakesmith  (Oxford University, UK) |  |
| Experimental model: Cell lines |  |  |
| HK-2 cells | ATCC | CRL-22 |

**EXPERIMENTAL MODEL AND SUBJECT DETAILS**

**Human subjects**

The tissue and serum were collected under the protocol approved by the Clinical Research Ethics Committee of the University of Florida (IRB201601019) following written informed consent from subjects in accordance with the Declaration of Helsinki Principles^2^.

**Mice**

All experiments were performed following the National Institutes of Health and Institutional Animal Care and Use Guidelines and were approved by The Animal Care and Use Committee of the University of Florida. The mice were maintained at 23°C (range ± 2°C), 30-70% humidity, and a 14:10-h light: dark cycle. Hepcidin knockout mice (*Hamp^-/-^*) were obtained from Sophie Vaulont (Institut Cochin, France), and their wild-type littermate controls were housed and maintained in the animal facilities of the University of Florida on a regular diet. 10-12-week-old male mice were used in these studies. The genetic *Hamp^-/-^* mice are characterized by plasma iron overload, high transferrin saturation, multi-visceral iron accumulation, and ferritin buildup^3,4^. We also utilized the inducible *Hamp^-/-^* (*iHamp^-/-^*) knockout mice obtained from Alexander Drakesmith (Oxford University, UK). In these mice, *Hamp1* alleles are floxed. They can be conditionally excised at any age by activating a ubiquitously expressed Cre-recombinase fused to a mutated estrogen receptor ligand-binding domain (CreERT2)^5^.

**Fungal strain and mouse model of systemic candidiasis**

The Candida albicans strains SC5314 was purchased from ATCC. *C. albicans* candidalysin deficient strain *ECE1 KO (ece1ΔΔ)* and its isogenic candidalysin sufficient strain *ece1ΔΔ+ECE1*(*ece1Δ* revertant, with comparable virulence to SC5314)^6^ were a kind gift from Dr. Bernhard Hube, Hans Knoell Insitute, Germany. The description of the red fluorescent C*andida albicans* strain CAF2-1-dTomato used in this study is provided in previous publications^1^. All the strains were streaked on agar plates composed of yeast extract, peptone, and dextrose medium containing penicillin and streptomycin (Gibco) at 33°C. Single colonies were added to the broth of identical composition and grown in a shaking incubator at 33°C. Cells were centrifuged, washed in PBS, counted using a hemocytometer, and injected into *Hamp^-/-^* or littermates (wild type) mice via the lateral tail vein. 2- 4X10^5^ *C. albicans* yeast cells were injected per mouse.

The iHamp*^-/-^* mice received 1 mg tamoxifen for three consecutive days (intraperitoneally). Following the tamoxifen regimen, these mice were infected with *C. albican*s (SC5314). In a cohort of iHamp*^-/-^* mice, following tamoxifen and *C. albicans*, were daily administered 50 nM mini-hepcidin PR-73 (a synthetic hepcidin agonist) intraperitoneally^7,8^ \ post tamoxifen and *C. albicans* injections. The first dose of PR-73 was 4 hrs post C.albicans injection. Experiments were terminated on day 3-4 post-infection.

**Induced liver fibrosis model**

8-10 weeks old C57Bl6 mice were used in this study. Carbon tetrachloride (CCl_4_) (Sigma) was injected to induce liver inflammation^9,10^. 0.3% CCl_4_ (diluted in olive oil) was injected intraperitoneally, twice weekly for 5 weeks at a dose of 10 uL/g of body weight. Two days after the last dose, animals were euthanized, and organs were harvested for further studies.

**Fungal Burden determination**

50 uL blood was collected by tail bleeding the mice 20 hrs post-infection, serially diluted in PBS, and 10 uL of each dilution was plated on yeast extract, peptone, and dextrose agar plates (YPD agar, Difco) containing penicillin and streptomycin. CFUs were determined after 24 hours of incubation at 37°C and results were expressed as CFUs/mL. The mice were euthanized on days 3 and 6 after infection to determine the tissue fungal burden in the kidney. The kidney, spleen, and liver were aseptically removed and homogenized using a tissue homogenizer. The tissue homogenates were serially diluted and CFU was measured as mentioned above and expressed as CFU/organ.

**Biochemical assays and tissue samples**

Before euthanasia, animals were anesthetized with ketamine (120 mg/kg)/ xylazine (12 mg/kg) and blood was drawn from the axilla. All the tissue slices were fixed with 10% neutral-buffered formalin for paraffin embedding, and with periodate-lysine-paraformaldehyde fixative (PLP) to be frozen in optimal cutting temperature compound, or snap frozen in liquid nitrogen for subsequent RNA and protein extraction, and immunofluorescence.

**Histopathology analysis**

**PAS and H&E staining**

Kidney tissue was fixed in 10% formalin and was submitted to the molecular pathology core at the University of Florida. Five μM thick paraffin embedded sections were cut and stained for Periodic Schiffs (PAS), and hematoxylin and eosin (H&E) by the molecular pathology core at the University of Florida.

**Grocott Methenamine Silver Staining**

10% formalin-fixed tissue sections were deparaffinized in xylene and stained by Grocott’s methenamine silver staining to determine the morphological changes of *C. albicans* during infection in mice. The tissue sections were collected post-infection and processed. The protocol described by the manufacturer, Polysciences (described in the table) was followed for the staining.

**Perls Staining**

As described in our previous studies, 10% formalin-fixed tissue sections were deparaffinized in xylene and stained for Perl`s detectable iron deposits^4^. After washing excess reagent, tissue was counter-stained with nuclear fast red and imaged for blue iron deposits.

**Plasma Creatinine assay**

Serum and plasma creatinine was measured using a commercial assay as described by the manufacturer (Diazyme, description in resources table).

**Immunofluorescence**

Three-micron, PLP-fixed kidney sections were used for the immunofluorescence detection of lotus tetragonolobus lectin (LTL) positive proximal tubular epithelial cells. Briefly, tissue sections were air-dried and incubated with 0.3% Triton X100/ 10% horse serum in PBS for 30 minutes. After washing the sections with PBS, an anti-CD16/32 antibody was added to block F_C_ receptors. This was followed by incubation for 90 minutes with FITC-labeled LTL (Vector Labs, 1:300). The sections were then washed 3 times in PBS and mounted with ProLong Gold antifade agent with (Life Technologies). The sections were imaged on a Keyence BZ-X800 fluorescence imaging microscope.

**In vitro detection of exposed β-1,3-glucan:**

For exposed β-1,3-glucan staining, *ece1ΔΔ+ECE1* and *ece1ΔΔ Candida albicans* were grown in YPD broth at 33°C 200 RPM for 16 hours. After washing with PBS, an aliquot was resuspended in YPD broth with or without 100 μM FeCl_3_ and grown overnight. Two million yeast cells from each condition were added on a 15 μM thick glass coverslip in a 12-well plate at 37°C. After overnight growth, the coverslips were fixed in 4% paraformaldehyde for 20 mins and sequentially treated with 0.5% Triton X-100 for 5 minutes, ice-cold 3% BSA/PBS for 1 hour and primary antibody for β-1,3-glucan (Mouse monoclonal) for 90 minutes (1:800 in 3% BSA-PBS). This was followed by incubation with Goat anti-Mouse PE, (1:600) and FITC conjugated anti-Concavilin A (30 μg/mL) for 45 minutes in the dark. After PBS wash, the coverslips were mounted with ProLong mountant without DAPI (Life technologies). Sections were imaged on a Keyence BZ-X800 fluorescence imaging microscope.

**In vivo detection and quantification of exposed β-1,3-glucan:**

WT and *Hamp^-/-^* mice (n = 5) were infected (intravenous) with two hundered thousand red fluorescent *Candida albicans* (CAF2-1-dTomato)^1^ and the kidneys were harvested 3 days later. An entire kidney was homogenized (Tissue Tearor, BioSpec Inc) in 1 mL PBS for 20-30 seconds. The homogenate was filtered through 100 μM filter and fixed in 4% PFA for 20 mins. Subsequent steps and antibody dilutions were as described above. Goat anti-Mouse Alexa fluor 647 (1:600) used to detect β-1,3-glucan. After PBS wash, the digest was diluted in 300 μL ProLong mountant without DAPI (Life technologies) and after vortexing, 100 μL was added onto a glass slide (Fisher, Superforst Plus) and mounted with cover slips. 4 random, 20X images of each slide were taken on a Keyence BZ-X800 fluorescence imaging microscope. Red hyphae with exposed β-1,3-glucan (green) colocalized as yellow were counted.

**Flow Cytometry**

Single-cell suspensions from the kidney were prepared as described in our previous publication^11^. The kidney was cut into small pieces and digested with collagenase (type 4; Worthington) and 100 μg/mL DNAse (Roche) for 20 minutes at 37°C. The digested kidney was then passed serially through a 70 μm and 40 μm sieve to collect the cell suspension. The cells were then incubated with eBioscience^TM^ Fixable Viability Dye eFlour^TM^ 780 (Invitrogen) (1:3000) and anti-CD16/32 (Fc block, clone 93; BioLegend, San Diego, CA) (1:100) in PBS for 20 mins in the dark. After washing with FAC buffer, cells were stained with PE-Cy7 conjugated anti-CD45 (30-F11), FITC-conjugated anti-CD11b (M1/70), PE-conjugated anti-Ly6c (HK 1.4) APC-conjugated anti-Ly6G (1A8 BD Bioscience). Flow cytometry data were acquired using Cytek Arora 3 laser (Fremont, CA). 500,000 events/samples were acquired and analyzed with FlowJo software 9.0 (Tree Star Inc., Ashland, OR).

**Real-time PCR**

For RNA isolation, frozen tissues were re-suspended in RLT buffer (Qiagen Inc., Valencia, CA) and homogenized using the TissueLyser system (Qiagen). For isolating fungal RNA, infected kidneys were homogenized in 2 mL Trizol and bead-beaten using zirconia beads for 45 seconds twice (with 5-minute intervals on ice). After spinning for 2 mins at 12,000 rpm the supernatant was added to 200 microliters of chloroform and phase separated. The top RNA layer was isolated and further purified using a RNeasy Plus mini kit (Qiagen). After loading RNA on the spin column (Qiagen), on-column DNA digestion (Qiagen) was performed for 30 mins at room temperature. Total RNA from tissue homogenates was then purified using the RNeasy Plus mini kit (Qiagen) following the manufacturer’s instructions. 1 μg of RNA was used to synthesize cDNA using the iScript cDNA synthesis kit (Bio-Rad Laboratories, Hercules, CA). The cDNA template was mixed with iTAQ SYBR green universal super mix (Bio-Rad) and quantitative PCR was carried out on a CFX Connect system (Bio-Rad). Data are expressed as fold change over control and were calculated using the 2^-△C(T)^ method. PPIA was used as the reference gene for mice and TDH3 for *C. albicans*. All the primers used in this study and their sequences are listed in the resource table.

**In-vitro studies**

**Cell line**

HK-2 cells (ATCC, CRL-2190), a human proximal tubular epithelial cell line, were maintained in ATCC-recommended Keratinocyte Serum-Free Medium (Gibco) supplemented with Bovine Pituitary Extract (0.05 mg/mL) and Human Recombinant Epidermal Growth Factor (5ng/mL) at 37C and 5% CO­_2_.

**Influence of iron on the growth of *C. albicans* in mouse kidneys and human PTEC cell lysates.**

The kidneys of C57BLK/6 mice were excised, cut into small pieces, and digested with 1.2 mg/mL collagenase (type 4; Worthington) and 100 μg/mL DNAse (Roche) for 20 minutes at 37^O^ C. After washing, the whole kidney digest was cultured for 48 hours in renal epithelial cell growth basal medium 2 (PromoCell, Heidelberg, Germany) supplemented with recombinant human epidermal growth factor (10 ng/ml), recombinant human insulin (5 μg/ml) epinephrine (0.5 μg/ml), hydrocortisone (36 ng/ml), human holotransferrin (5 μg/ml), triiodo-L thyronine (4 pg/ml), and 0.5% fetal calf serum (28515173). After 24hrs, 50 or 100 μM ferric ammonium citrate was added to the cultures. Forty eight hrs later, the adherent cells were washed, treated with trypsin and lysed in water. The supernatant were bought upto 1X PBS and spiked with 10,000 *C. albicans* yeast cells and growth curves were generated for 19 hrs at 32^O^ C.

To evalaute the growth of *C. albicans,* HK-2 cells (2X10^5^/well) were iron overloaded using Ferric Chloride (100 µM) as described by van Raaji et al. (30806852). After washing to remove any non-internalized iron, the vehicle or iron-loaded HK-2 cells were treated with *C. albicans* (50,000/well) for 20 hrs. The supernatant was inoculated with 10,000 C. albicans, and growth curves were generated for 12 hours at 32^O^ C.

**Effect of iron on *C. albicans* hyphal sustenance.**

To determine the role of iron in hyphal sustenance, *C. albicans ece1ΔΔ+ ECE1* and *ece1ΔΔ* were grown for 18 hours in YNB broth supplemented with 2% glucose, 5 gm/L NH4SO4, 0.79 gm/L amino acid supplement, and 2 mM/L CuSO4 at 33^O^ C. After 18 hrs, an aliquot was resuspended in YNB broth supplemented with ammonium sulfate, copper sulfate with or without 100 mM ferric ammonium citrate (source of iron) as described by Tripathi et. al (PMID:32503842). The next day, 1 X 10^6^ yeast cells were plated in 12 well plates in the same broth and grown at 37^O^ C. Images were taken at 0 hours, 3 hours, and 24 hours using Keyence BZ-X800 microscope*.*

**Effect of *ece1ΔΔ* *C. albicans* grown in excess iron on human proximal renal tubular cells immune response.**

The *ece1ΔΔ Candida albicans* were grown in YPD broth at 33°C 200 RPM for 16 hours and an aliquot was resuspended in YPD broth with or without 100 μM FeCl_3_ for overnight growth. After washing, the yeast cells were resuspended in keratinocyte-SFM media and added to HK-2 cells (MOI 1:3 Cells: Fungus) at 37^0^C. Supernatent were collected at different time points. IL-8 was measured using a commercial sandwich ELISA as recommended by the manufacturer. *ece1ΔΔ C. albicans* was grown in standard and iron-rich broth (as above) and then added to HK-2 medium, to follow their growth at 37^0^ C for 4 hrs.
